# LVV SMRTcap reveals extensive proviral variation in lentiviral vector-transduced CAR T cells

**DOI:** 10.64898/2026.05.13.724601

**Authors:** Catherine W. Kaiser, Ghazal S. Mehs, Erin M. Elliott, Joanna E. Mroczkowska, Ankita Jain, Michael Ferguson, Alfred L. Garfall, Frederic D. Bushman, Joseph A. Fraietta, Eric C. Rouchka, Melissa L. Smith

## Abstract

Lentiviral vectors are commonly used to introduce chimeric antigen receptor transgenes into T cells, but routine assays quantify vector copy number or integration sites without sequencing full-length integrated vectors. HIV-1 proviruses often acquire large deletions and cytidine deaminase-driven hypermutation; whether similar variation occurs in therapeutic lentiviral vectors is unclear. We adapted a novel long-read capture approach to enrich long fragments spanning vector DNA and adjacent human sequence, enabling simultaneous integration-site mapping and proviral integrity analysis with single-molecule resolution. In research-grade CAR T cells produced with an experimental, transient-transfection lentiviral vector workflow, 40% of integrated vectors carried recurrent deletions that removed the internal promoter or parts of the chimeric antigen receptor cassette. The dominant promoter deletion was present in the viral stock. In clinical chimeric antigen receptor T cell products, promoter deletions were less frequent, but detectable pre-infusion and post-infusion. Across datasets we observed widespread G-to-A substitutions consistent with restriction factor editing, including changes predicted to introduce premature stop codons within the transgene open reading frame. Our method reveals proviral variants invisible to standard quality-control assays and provides a framework to improve vector production and monitor transgene integrity in clinical products.

## Introduction

Lentiviral vectors (LVVs) are human immunodeficiency virus (HIV)-derived gene therapeutics that have been extensively modified from the original viral genome to deliver therapeutic transgenes and block ongoing vector replication. There have been four increasingly modified generations of LVV systems, with the third-generation vectors being the most prominently studied and used in the clinic (1–3). In cancer immunotherapy, LVVs commonly deliver a chimeric antigen receptor (CAR) transgene to T cells, generating CAR T cells directed against a tumor-associated antigen (2). CAR T cell therapy can induce durable remissions, particularly in B-cell malignancies, yet responses vary substantially across patients and products (4, 5). Five of the seven Food and Drug Administration (FDA)-approved CAR T cell therapies are LVV-derived; however, in November 2023, all clinical CAR T cell therapies were placed under federal investigation due unexpectedly high rates of CAR(+) malignancies leading to adverse medical events including death (6). Thus, because LVV-mediated gene transfer is the defining modification in most approved CAR T therapies, understanding the integrity of the integrated vector in the final cell product is important for both efficacy and safety.

After entry into a target cell, the single-stranded RNA genome packaged in the LVV particle is reverse transcribed into double-stranded cDNA, which then integrates into the host genome. LVV integration favors transcriptionally active genomic regions and is enriched in introns (7). Another unique feature of integrated HIV is the acquisition of single nucleotide variants (SNVs) and large internal deletions upon reverse transcription and integration. This maintenance of “proviral integrity” has been less explored in the context of LVV-derived gene therapies (7–10). The enrichment of HIV SNVs can be primarily attributed to the high error rate of the HIV reverse transcriptase and restriction factor editing during virion production and maturation. Internal deletions within the HIV provirus are thought to arise from looping of the viral RNA during the RT process. Characterization of the latent HIV reservoir by multiple groups has demonstrated that at least 90% of proviruses contain large internal deletions (11, 12). The apolipoprotein B mRNA-editing enzyme catalytic polypeptide-like 3 (APOBEC3) family of cytidine deaminases have evolved to inhibit intracellular RNA virus replication. APOBEC3 family genes induce C-to-U mutations in the ssRNA genome resulting in G-to-A mutational signatures in the ds-cDNA later integrated into the host genome (13, 14). HIV has evolved to combat APOBEC3-mediated editing through the evolution of an accessory gene, “virion infectivity factor” (*vif*), which targets APOBEC for degradation, protecting HIV-1 genome integrity (15). *Vif* was removed from the third generation LVV systems widely used today to increase vector safety and viral protein-associated inflammation (16–18). Hart et al. has only recently identified APOBEC3-related hypermutation signatures in integrated LVV cassettes (10). Limited work has been done thus far to investigate whether hypermutation and large structural variations are consistently present in therapeutic LVV systems.

Prior studies have investigated a possible link between LVV integration sites and genotoxicity and/or transformation in CAR T cells (19–21). Although integration of CAR-delivering LVVs has been linked to oncogene expression, it occurs rarely (21, 22). LVV integration sites are typically monitored using ligation-mediated PCR (LM-PCR) followed by short-read, high throughput sequencing, while transgene abundance (e.g., CAR) is evaluated using vector copy number assays via qPCR to quantify transgene levels once cells have been transfused back into the patient (23, 24). Neither of these assays evaluate the full integrated LVV cassette with nucleotide level resolution, nor are they capable of resolving large insertions/deletions (indels). Moreover, the use of short-read NGS to resolve integration sites may not uniquely map integration events within repetitive elements, which make up ∼50% of the human genome (25).

To address this gap, we adapted our single-molecule, real-time capture assay (SMRTcap) to enrich long genomic DNA fragments spanning LVV sequences together with adjacent human flanks (26). This approach (“LVV SMRTcap”) enables simultaneous, single-molecule resolution of: (i) LVV integration sites, (ii) clonal expansion inferred from shared integration sites and distinct shear points, and (iii) proviral integrity across much of the integrated cassette (26). Applying LVV SMRTcap to research-grade CAR T cells and to clinical pre- and post-infusion CAR T samples, we identify recurrent internal deletions that can remove the internal promoter or parts of the CAR cassette, and we detect widespread G-to-A substitutions consistent with APOBEC3 editing, including variants predicted to introduce premature stop codons in the CAR open reading frame.

## Results

### LVV SMRTcap captures integration site distributions comparable to LM-PCR

Recently we published HIV SMRTcap, designed for comprehensive characterization of the HIV-1 reservoir (26). To adapt and validate the SMRTcap pipeline (Figure 1A) for LVVs, we produced research-grade CAR T cells by transducing four peripheral blood mononuclear cell (PBMC) samples from unique healthy donors with an LVV vector carrying a CAR in development (Figure S1). We then performed side-by-side integration site validation using LVV SMRTcap and LM-PCR, according to the Serrao, et. al., protocol (Figure 1B) (27). Since LVV SMRTcap and LM-PCR were performed on separate genomic DNA aliquots from the same donor-derived cell populations, we focused on concordance of global integration patterns rather than on exact site-by-site overlap (Figure 1C); matched data for the remaining three samples are presented in the Supplemental files (Figure S2).

**Figure 1:**
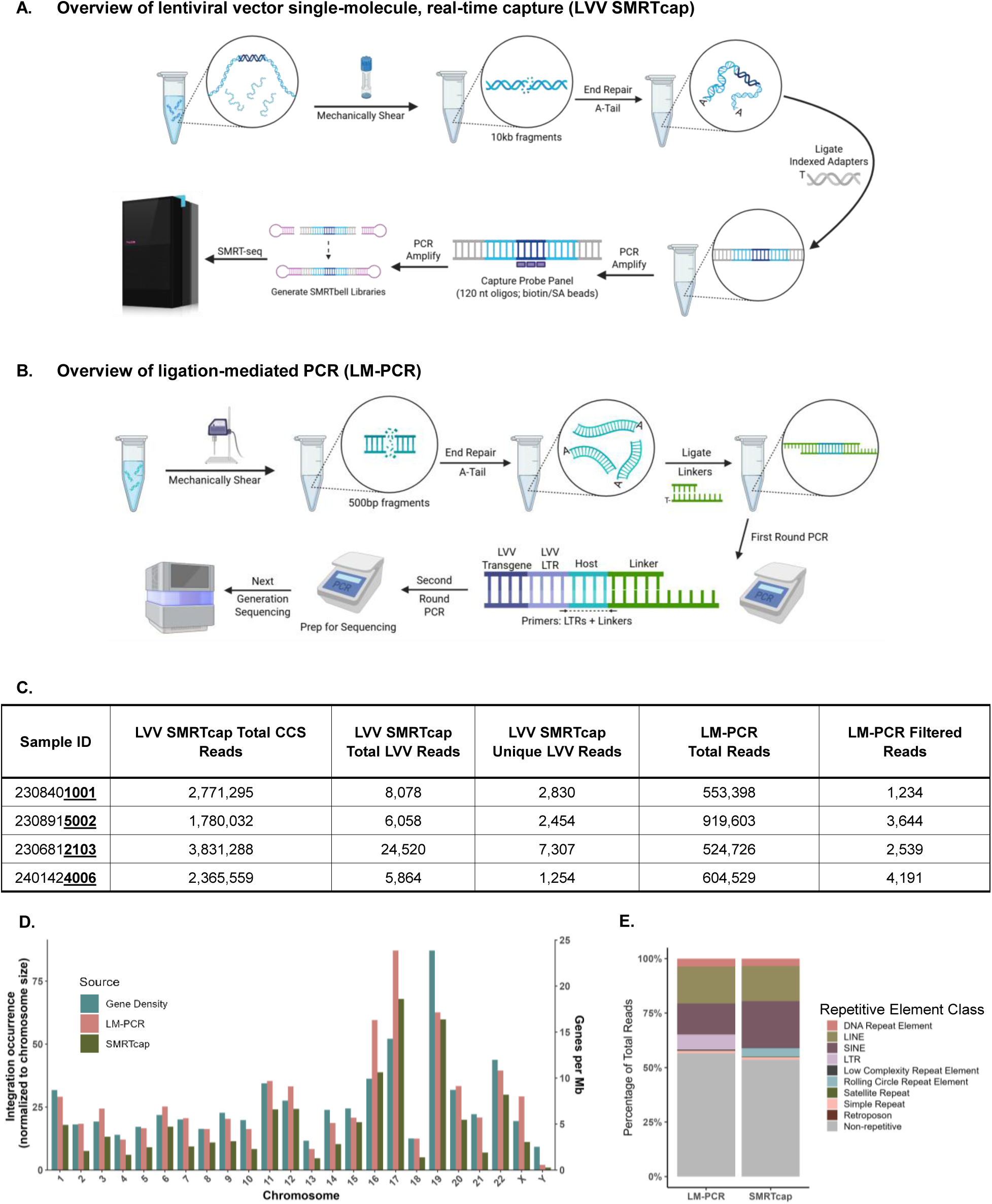
LVV SMRTcap and LM-PCR integration profiles are concordant in distribution and frequency. **A)** Overview of LVV SMRTcap pipeline: LVV-containing gDNA is mechanically sheared to 10-20kb and gDNA fragments are end repaired, A-tailed, and barcoded adapters are ligated to allow for non-specific, bulk amplification. Biotinylated oligonucleotide probes specific for the integrated LVV genome target are used to enrich DNA fragments containing proviral DNA and streptavidin beads are used to isolate fragments containing viral template bound to biotinylated probes. Post capture templates are amplified with a second round of PCR and undergo SMRTbell template preparation. All samples were sequenced using highly accurate single-molecule sequencing on the Revio system. Created using BioRender **B)** Overview of LM-PCR method: gDNA is sheared to ∼500bp, end repaired, A-tailed, and linkers are ligated. Linkers allow for unbiased mass amplification, and primers are used for targeted first-round amplification of fragments with lentiviral LTR-containing ends. Primers used in the second amplification step allow for annealing of sequencing adapters. Libraries are sequenced as 2×250 on a MiSeq V3 flow cell. Created using BioRender **C)** Research-derived CAR T cell donors and read metrics generated by LVV SMRTcap and LM-PCR. **D)** LVV SMRTcap and LM-PCR viral integration coverage per chromosome from donor 2103, normalized to chromosome size. Gene density is presented using Cold Spring Harbor Laboratory Press’s Guide to the Human Genome, genes per sequenced Mb are recorded for each chromosome (48). **E)** LVV SMRTcap and LM-PCR integration site composition into unique and repetitive elements from representative donor 2103.

While all four donor-derived CAR T cell lines were processed by both LVV SMRTcap and LM-PCR, each assay was performed with unique genomic DNA inputs. Due to this sampling difference, we do not anticipate exact concordance across both methods but rather compared the overall distribution and frequency of integration patterns observed (Figure 1D, S2). For example, integration rates into chr17 and chr19 are relatively higher per kilobase using both LVV SMRTcap and LM-PCR, while chr13 and chr18 show relatively fewer integrations per kilobase. Further, as has been previously shown, integration frequencies detected by both methods were associated with gene density per chromosome, providing further validation of the LVV SMRTcap pipeline results align with expected outcomes (28). We anticipated that, due to longer reads in the host flanking regions improving mappability, LVV SMRTcap may have increased resolution identifying integration events in repetitive genomic elements. This may be the case for integration in SINEs (Figure 1E, S3), as 14% total integrations were identified in SINEs by LM-PCR, while 21% were identified by LVV SMRTcap (Figure 1E), although we cannot discount that this increase in LVV SMRTcap was due assay-specific sampling. Lastly, while LVV integration occurred in many types of genomic elements, most integration was found within intronic regions, as previously described (Figure S4). Together, these data validate the ability of LVV SMRTcap to accurately resolve LVV integration patterns to the specificity and extent of commonly used LM-PCR based methods.

### LVV SMRTcap reveals recurrent internal deletions in research-grade CAR T cells

Both the HIV SMRTcap and LVV SMRTcap protocols depend on non-specific mechanical shearing of genomic DNA (gDNA) to generate the longer (>8 kilobases, kb) fragments used as capture input. This random process can shear in the middle of an integrated provirus, resulting in fragments with only one flanking human DNA sequence (Figure 2A). For donor 2103, we observed that 52.7% of LVV SMRTcap sequence reads contained a single human flank (e.g., either on the 5’ or 3’ flank of the integrated provirus), 23.8% LVV fragments (e.g., contained no human flanks, potentially residual plasmid or episomal LVVs), and 23.5% were dually flanked (e.g, both 5’ and 3’ human flanks are present) (Figure 2B). Across all four donor samples, captured fragments ranged in size from 2–12 kb, with an average read length of ∼7 kb (Figure S5D–G). Given that the research LVV-CAR genome used is 6.1kb, based on this size distribution, it is not surprising that a substantial number of fragments contain only a single flank (Figure S5A-C). To visualize proviral integrity and investigate internal deletions, we started our analyses of proviral intactness with the subset of dually flanked reads, since we have complete confidence in the enriched proviral content between these flanks. First, the complete proviral insert sequences were mapped to the viral genome reference for integrity analysis. We observed a recurring 812bp deletion spanning the transgene promoter in a substantial proportion of the dually-flanked proviral reads (Figures 2C, S6-S8). We then extended our analyses to include single-flanked reads, provided this proviral region was retained during random shearing process. In total, this deletion was found in approximately 40% of all integrated provirus enriched from donor 2103. When examined, the other three donor CAR T cells showed very similar frequencies of this specific deletion spanning the CAR-driving promotor (Figure 2D). We also observed recurring structural variation unique to each donor (Figures 2C, S6-S8). For example, donor 2103 carried a 642-base deletion in the CAR gene cassette, which was in phase with the promoter deletion in ∼0.1% of inserts (Figure 2C), and a 171-base deletion within the 3’ LTR (0.22% of inserts).

**Figure 2:**
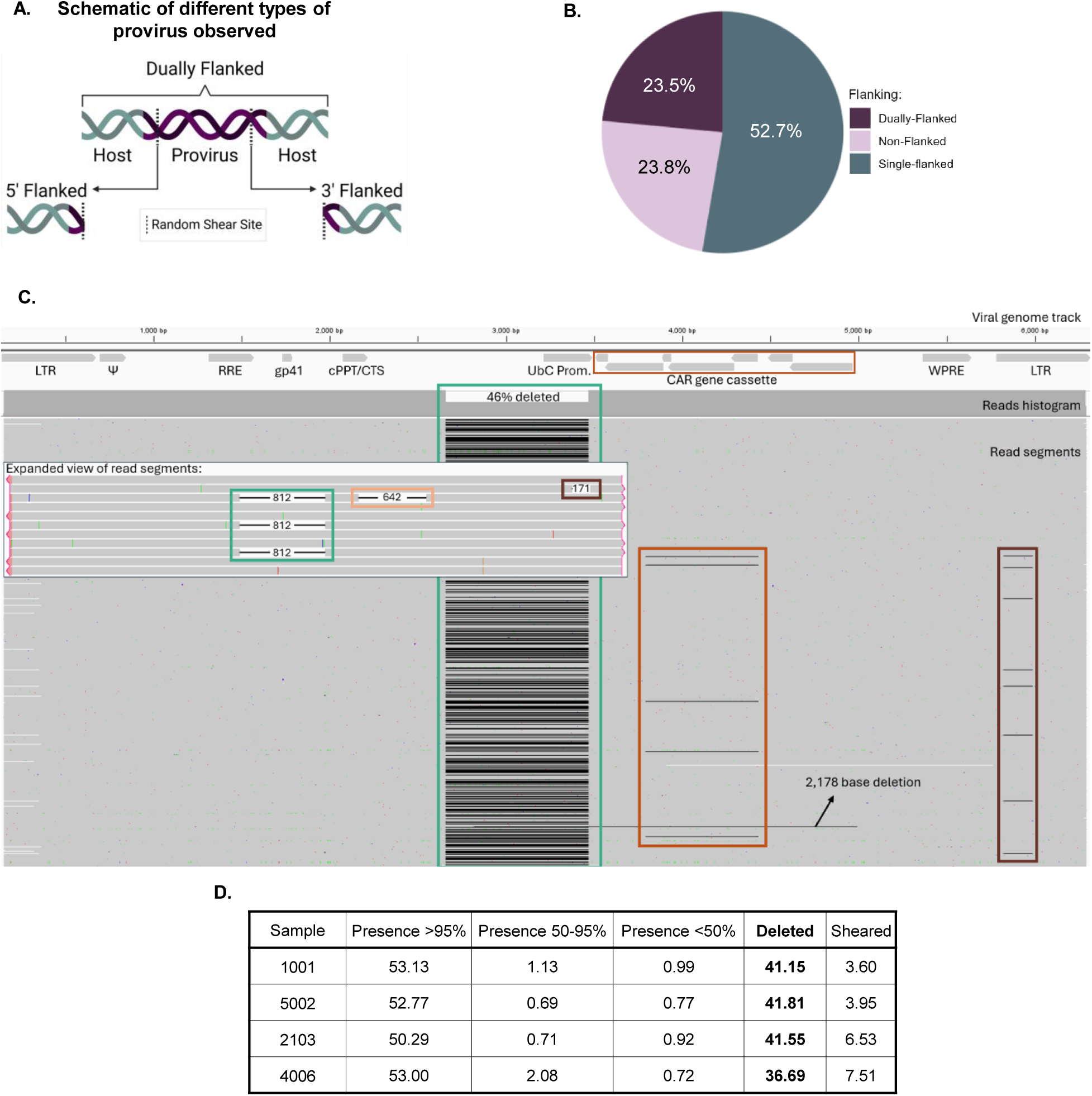
Large internal deletions characterized across research CAR T cell samples using LVV SMRTcap data. **A)** Overview of all sequencing output moieties derived from random shearing of proviral viral templates. Created using BioRender **B)** Viral shearing read length profiles for donor 2103 from LVV SMRTcap data. **C)** Dually flanked proviruses from donor 2103 mapped back to the viral genome reference and visualized using IGV. Viral genes are annotated beneath the viral genome track. Read segments include deletions are annotated as black lines. An expanded view of read segments shows recurring deletions with matching outlines to show their position in the unexpanded view. Percentage of dually flanked reads with 812 base deletion is annotated. **D)** Recorded integrity rate for the UbC promoter across research CAR T samples. ‘Sheared’ portion is due to random mechanical shearing prevents this portion from being analyzed in the sequence that was captured.

We wanted to ensure the promoter deletion was not an artifact of our in-lab CAR T cell generation system. First, we confirmed that the plasmid stock encoding the CAR-carrying LVV did not contain this internal deletion by sequencing the viral plasmid using Oxford Nanopore Technologies (ONT). The raw ONT reads (FASTQ) were mapped to the viral plasmid and the analysis showed an intact plasmid with no internal deletions detected even at low frequencies (Figure 3A). Next, we examined the infecting LVV stock generated by the 6-plasmid, fourth-generation LVV transfection system (see Materials and Methods). Lentiviral particles were purified over a 20% sucrose gradient, and viral mRNA was isolated and used as input into ds cDNA generation as part of long-read full-length isoform sequencing preparation (IsoSeq), followed by single-molecule sequencing on the Revio system. Sequencing of the purified viral stock revealed that the 812-base deletion spanning the human ubiquitin C (UbC) promoter (Figure 3B) was introduced at the viral production step. However, additional variation was found post-integration suggesting SNVs and insertions/deletions were acquired both during viral particle production and upon integration.

**Figure 3:**
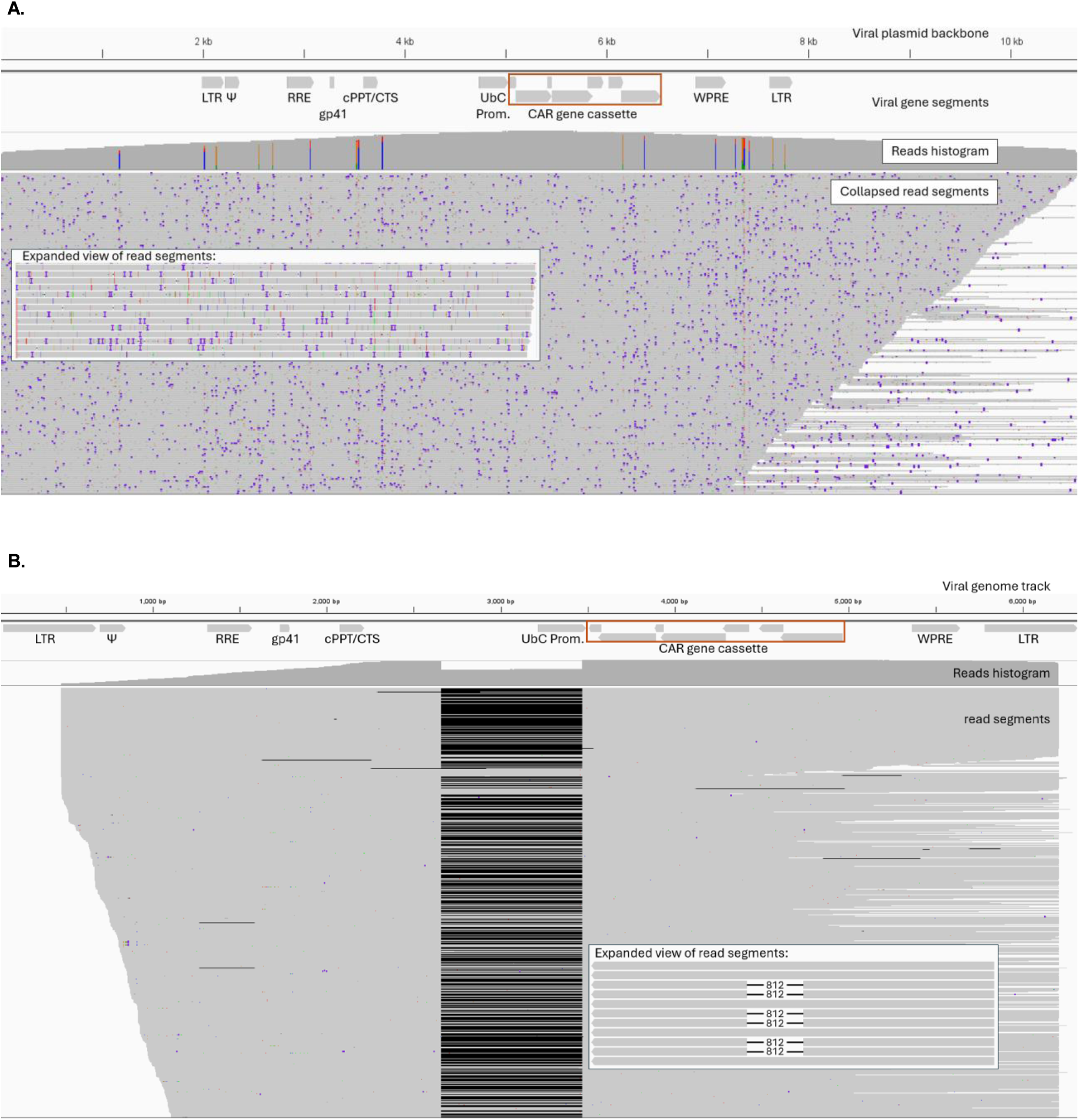
Internal viral genome deletions captured post viral production. Long read sequencing was used to evaluate, map and visualize viral data with IGV **A)** Plasmid vector was sequenced using Oxford Nanopore Technologies and mapped back to the viral plasmid reference. SNP and INDELs observed are technical errors introduced through nanopore sequencing. **B)** Sequencing of viral RNA from LVV stocks. Viral particles were purified, viral mRNA isolated, reverse transcribed and amplified, underwent SMRTbell library prep, and sequenced using the Revio system, followed by mapping back to the viral genome. Insertion events are depicted as purple hashes, showing insertions >2 bases in length. The read histograms are set to show SNPs that occur in >10% of reads. Black lines depict deletion events; expanded view of reads show the most recurrent deletions per sample.

### Clonal expansion can enrich proviral variants predicted to compromise CAR expression

Beyond cataloging integration events, LVV SMRTcap can infer clonal expansion by combining the integration site with the unique genomic shear points that define each captured DNA fragment (Figure 4A) (26, 29, 30). Using donor 2103 again as a representative sample, we noted that most clonal clusters were relatively small, including only 20-30 unique templates and that none of the donor samples were dominated by a single expanded clone (Figure 4B). This was not surprising, as these cells were only allowed to expand in culture for 48 hours before harvest. We extended our analysis to unique deletions captured in single-flanked reads that were clonally expanded, schematized in Figure 4C. In donor 2103, we identified several templates with unique indel patterns, including one clonal group (n=7 unique templates) with an insertion of 83 bases in the LTR that appears to be a result of a LTR duplication (Figure 4C). It is likely that we captured more expanded clones in donor 2103 due to the increased data generated; donors 1001, 5002, and 4006 did demonstrate unique deletion patterns in single-flanked reads but without demonstrable clonal expansion (Figures S6-S8).

**Figure 4:**
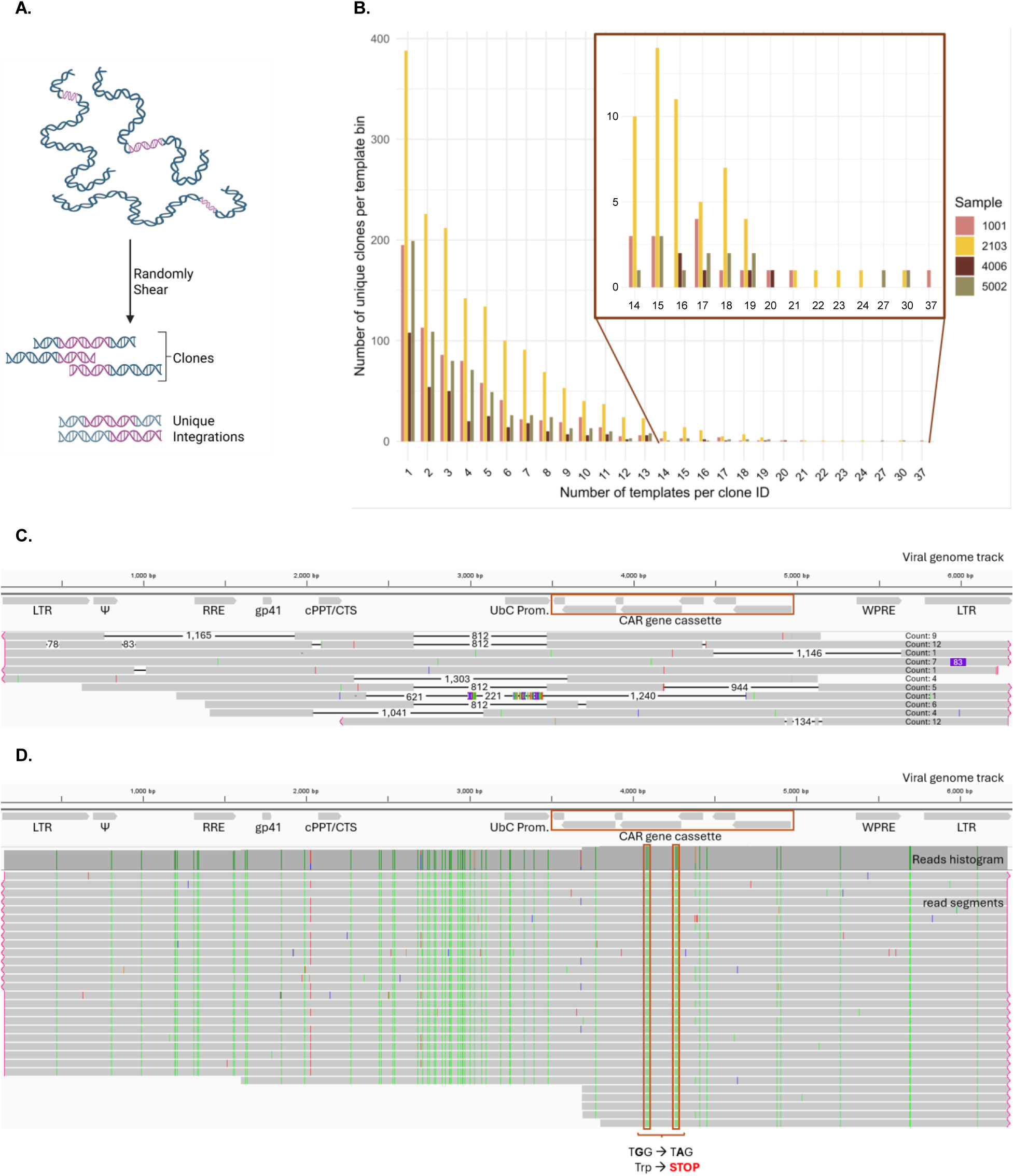
Clonal expansion of research CAR T cells shows structural variation. **A)** gDNA is randomly sheared, creating unique shear sites used as molecular identifiers for distinction of clones from unique integration events. Created with BioRender. **B)** Histogram of clonal expansion across research CAR T samples. The x-axis represents the number of templates for an individual clonal group, while the y-axis is the number of templates per clonal group; for example, x = 1 is a unique clone, whereas x = 3 is a clonal group containing 3 unique templates. **C)** Donor 2103 proviral insertions with unique deletion variants mapped back to the viral genome reference. Viral genes are annotated beneath the viral genome reference track and read counts on each read indicate clone size for each LVV insertion event. **D)** Clonally expanded LVV integration into *TMEM 131* from donor 2103, visualized as mapped to the viral genome mapped with IGV. Varying colors represent unique DNA bases, and highlight SNPs: A = green, G = orange, T = red, and C= blue. The read histogram shows SNPs occurring in >10% of reads.

When characterizing the most expanded clone in donor 2103 (unique template count of 20), we observed that, compared to the vector-encoding plasmid, the provirus demonstrated extensive G-to-A nucleotide conversions (Figure 4D). Furthermore, two G-to-A conversions within the CAR open reading frame resulted in putative premature stop codons, suggesting truncation of the CAR gene transcripts derived from this expanded clone. The CAR is known to be transcribed as a continuous transcript; therefore, these novel premature stop codons would inhibit expression of several critical CAR domains in this clone. We were able to return to the quality control profiling we had previously performed to investigate if these G-to-A mutations were found in viral plasmids or viral stock populations. We did not observe substantial G-to-A conversions in either the propagated plasmid or viral stock (Figures 3A and 3B). This indicates these modifications were made post-particle generation and during/or post-infection. The G-to-A patterns strongly suggest ABOBEC3-mediated editing, confirming a recent publication that observed similar patterns (10). Further, the largest clones observed in the CAR T cells generated from donors 1001, 5002, and 4006 all contained the 812-base deletion spanning the UbC promoter (Figure S9). Together, our clonal analyses suggest that, either because of hypermutation or structural variation, the largest expanded clonal group across all four donors likely expresses a non-functional CAR.

### Clinical CAR T cell products contain low-frequency promoter deletions and APOBEC-like mutational signatures

While LVV SMRTcap revealed extensive heterogeneity and hypermutation in our fourth-generation research-grade CAR T cells, we needed to assess if this would extend to clinical grade, third-generation, CAR T products. Collaborators at University of Pennsylvania provided access to pre- and post-infusion CAR T cells from recent clinical studies for LVV SMRTcap characterization. A third-generation lentiviral vector production system was leveraged to generate the LVV used to transduce these autologous patient T cells, resulting in CAR T cells targeting either B cell maturation antigen (BCMA, patients 46417-37, 14415-13 and 14415-15), or B cell marker CD19 (patient 19416-04).

We first characterized the pre-infusion products using LVV SMRTcap (Figure 5A). Integration frequency across the genome and into specific genomic features showed similar trends as in the research samples and previously published data (Figure S10). When examining proviral integrity, we observed recurring insertion and deletion patterns within the proviruses (Figure 5B, S11-S13). Data from all four pre-infusion samples showed a small subset (range 0.07%-1.09%, median 0.315%) of reads carrying a 939 bp deletion spanning the eukaryotic translation elongation factor 1 alpha (EF1a) promoter. A median of 0.54% of reads (across all four pre-infusion samples) contained some type of deletion in the EF1a promoter driving CAR expression (range: 0.35%-1.29%). Further, patient 19416-04 pre-infusion CAR T cells carried an in-frame 6 base insertion between the promoter and the CAR gene segments in all reads captured except for those that contained a different 1,150 bp deletion spanning the EF1a promoter. Small patient-specific deletions and SNP variants were also observed within the pre-infusion samples (Figure 5B, S11-S13). As seen in the research samples, extensive G-to-A conversion was observable and annotated in each pre-infusion sample (Figure 5C). In one case, a single G-to-A conversion resulted in a predicted stop codon that would likely result in a truncated CAR in that provirus (Figure 5C, track 37).

**Figure 5:**
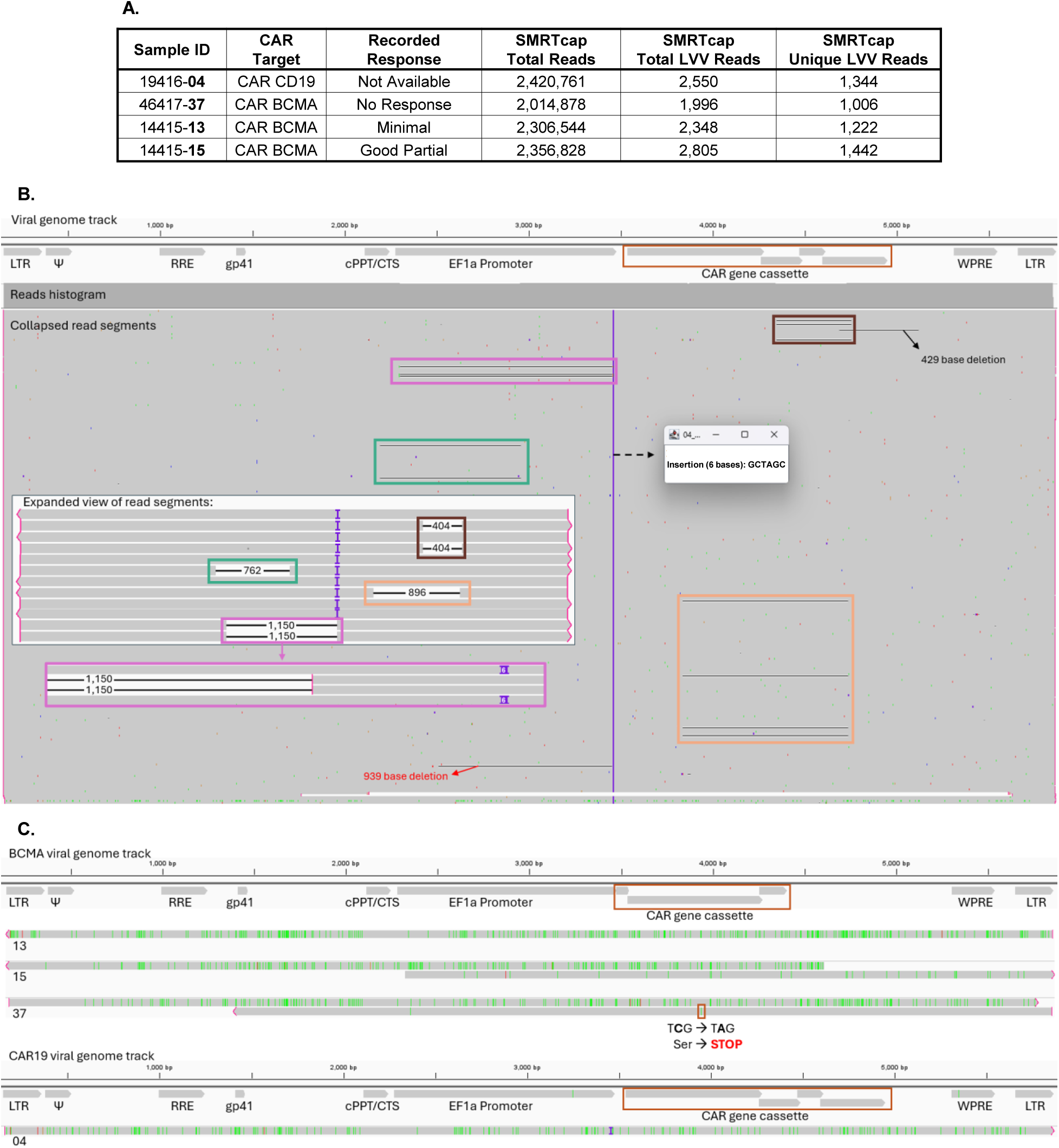
Clinically derived CAR T cell products contain structural variation. **A)** Patient details and LVV SMRTcap sequencing read metrics for all four characterized pre-infusion CAR T cell products. **(B)** Dually flanked viral insertion reads from representative donor 19416-04 mapped back to the viral genome reference. Blue blocks indicate insertion events >2bp. Black lines are deletion events in individual sequencing reads. The expanded view of read segments shows recurrent deletion patterns across the pre-infusion CAR T sample with colored outlines to indicate where they are occurring across the dually flanked reads. **C)** Example pre-infusion CAR T viral insertion reads from clinical donors with elevated evidence of G-to-A conversion events (shown in green) mapped back to the viral genome reference.

### Longitudinal post-infusion samples capture proviral variants in circulating cells

The impact of CAR T therapies occurs post-infusion and is based on which CAR T cells are activated and exert their desired effect. To evaluate whether those CAR T cells characterized as carrying hypermutated or deleted genomes pre-infusion could be observed during the treatment phase, we performed LVV SMRTcap on four longitudinal PBMC samples collected on days 7, 14, 21, and 28 post-infusion from patient 14415-15. As expected, LVV-containing reads were much less frequent in post-infusion PBMCs than in the pre-infusion products, reflecting dilution by non-transduced circulating cells and limited sampling of tissue-resident CAR T cells (Figure 6A). Despite this limitation, integration sites recovered from post-infusion samples were predominantly intronic (Figure 6B) and included clonally expanded groups at multiple time points, consistent with *in vivo* proliferation of some CAR T clones (Figure 6C).

**Figure 6:**
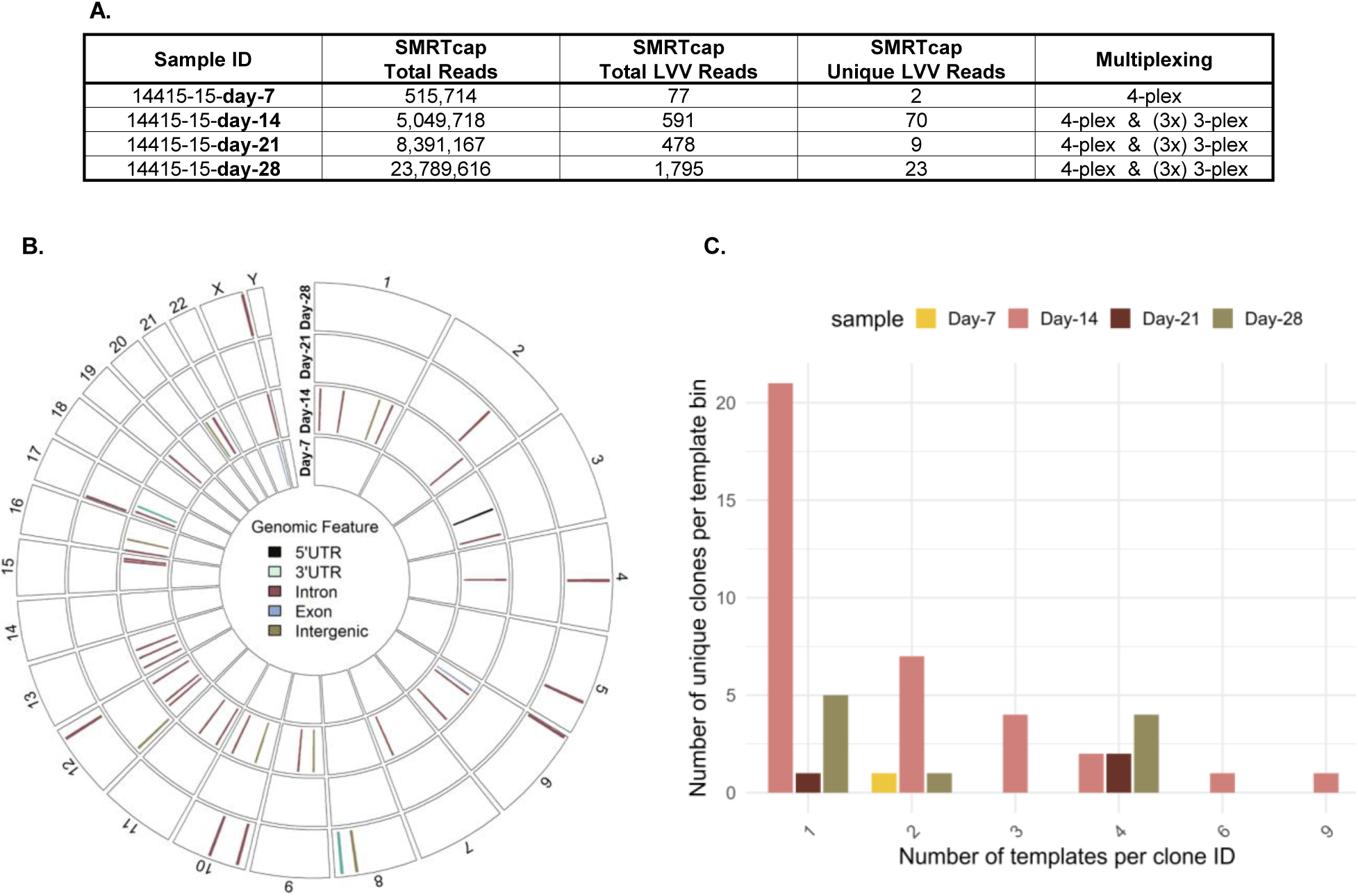
LVV SMRTcap profiling of longitudinal samples collected post-infusion from patient 14415-15. **A)** Post-infusion sample details and LVV SMRTcap read metrics. **B)** Chromosomal integration site coverage across each time point. Circos plot is divided amongst the chromosomes of the human genome, where each segment is a different chromosome. Each colored hash mark represents a unique recorded integration position along the chromosome. The color of the line used for marking the integration site corresponds to the genomic feature for that integration event (in integrity legend). **C)** Histogram of clonal expansion across post-transfusion longitudinal samples. The x-axis represents the number of templates for an individual clonal group, while the y-axis is the number of templates per clonal group; for example, x = 1 is a unique clone, whereas x = 3 is a clonal group containing 3 unique templates.

Within the limited number of proviral sequences recovered from these post-infusion samples, we observed both indels and SNVs (Figure 7). On day 7, we detected a provirus with a small deletion in the EF1α promoter, and a G-to-A substitution predicted to introduce a premature stop codon within the CAR cassette (Figure 7A). On day 14, most captured proviruses were intact across the sequenced region, although we observed a deletion spanning the central polypurine tract/central termination sequence and the start of the EF1α promoter (Figure 7B). The functional implications of such a deletion remain unknown. On day 21, we recovered a provirus with extensive G-to-A substitutions but without predicted stop codons in the CAR open reading frame (Figure 7C). On day 28, we again observed a G-to-A substitution predicted to introduce a premature stop codon within the CAR cassette (Figure 7D). Together, these data demonstrate that LVV SMRTcap can recover proviral variants from post-infusion samples, although deeper sampling will be required to quantify their prevalence *in vivo*.

**Figure 7:**
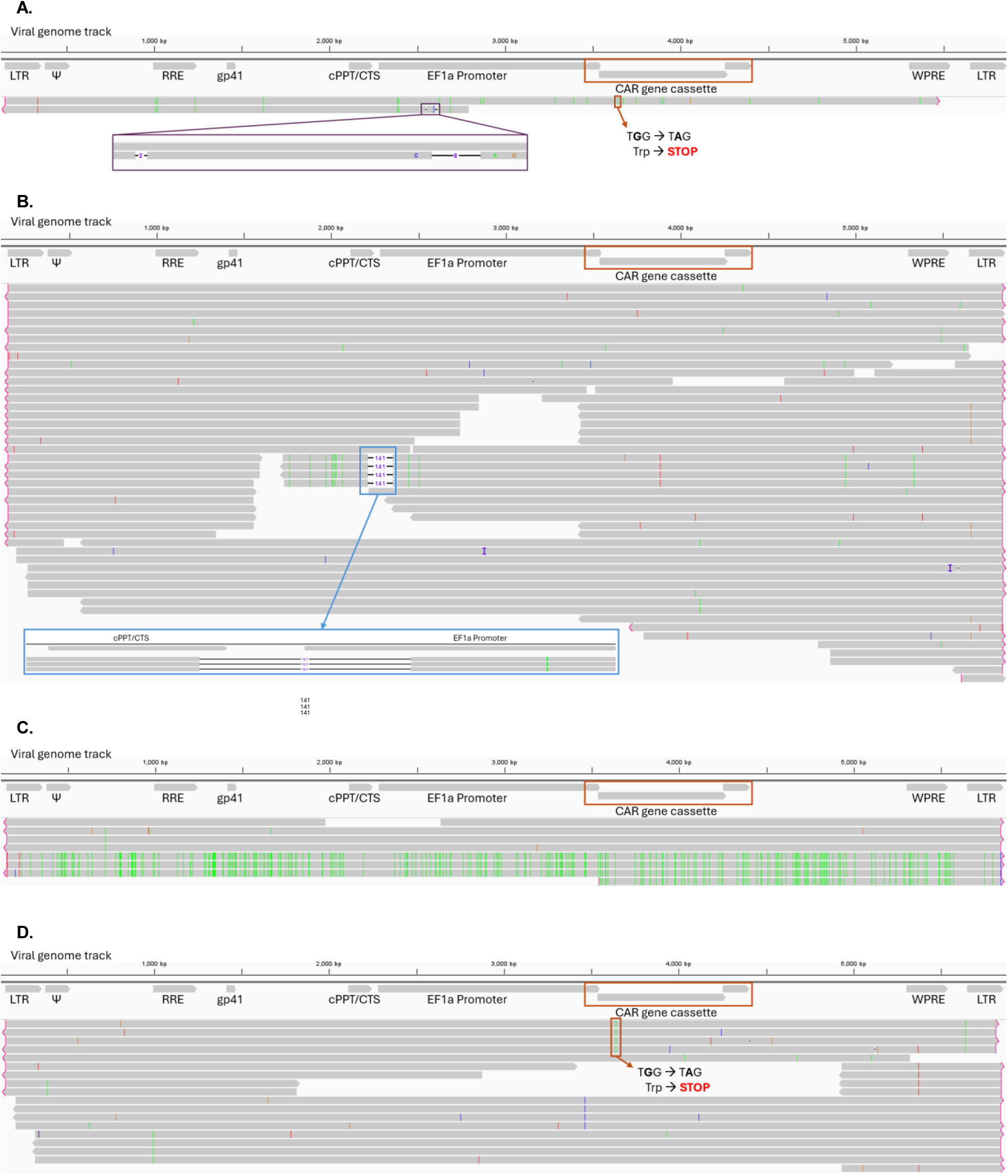
Characterized SNV and indels in patient 14415-15 CAR T cell samples provirus using LVV SMRTcap. Post treatment CAR T cell samples were taken at days: **A)** 7, **B)** 14, **C)** 21, and **D)** 28. Colors represent unique bases and highlight SNPs: A = green, G = orange, T = red, and C= blue. Reads histogram shows SNPs occurring in >10% of reads. Lines connecting read segments are used to depict deletions spanning that region, with the number present corresponding to the size of the deletion event.

## Discussion

LVVs are known to integrate into regions of active transcription, and in some rare cases, integration can drive increased CAR T cell proliferation leading to either recovery (31) or toxicity (21). LVV integration patterns and these few, unique cases have been extensively characterized; however, little attention has focused on understanding whether the LVV remains intact within therapeutic products. In clinical settings, vector copy number is often used to confirm successful engraftment by quantifying transgene copies by qPCR in patient-derived tissue samples. Depending on size of the amplified product and where the qPCR primers lie, this assay could also detect the presence of truncated LVV transcripts, if present. Integration site analysis using LM-PCR methods is typically only applied if there is suspicion of an adverse event, such as malignancy. Neither qPCR nor LM-PCR resolve proviral accumulation of SNVs or structural variation.

To address these limitations, we have adapted our recently developed HIV-SMRTcap pipeline (26) to resolve both LVV integration and proviral integrity in multiple sources of CAR T cells. We found that up to 40% of research-derived CAR T cells generated using a fourth-generation LVV contained large internal deletions spanning the UbC promoter driving therapeutic CAR expression. Upon further investigation, we determined this deletion was introduced during viral particle production and before host cell transduction, while several other SNVs and indels appeared to be integration related, as they were not present in viral stocks. Although not to the same extent, both SNVs and indels were observed in the pre-infusion CAR T cell products generated for clinical use. Though the post-infusion analyses were limited to only one patient, we did demonstrate that hypermutated and internally deleted CAR T cells survive, expand, and persist *in vivo.* In fact, predicted premature stop codons due to suspected APOBEC3-mediated mutation were found in the proviruses characterized from days 7 and 28 post CAR T cell treatment. Together, these results raise the question of what frequency of post-infusion samples contain these variations and if these mutations have any effect on cellular fitness or therapeutic efficacy.

We also observed extensive G-to-A conversion across almost all samples examined with LVV SMRTcap, suggesting APOBEC3 cytidine deamination activity was a major player impacting the extent of CAR T cassette polymorphism. In the data, G-to-A conversions often resulted in predicted premature stop codons in CAR T cell products. This was most notable in a post-infusion expanded clone at Day 21 from patient 14415-15. The extent of these mutations is not surprising as the APOBEC3 gene family has evolved as a natural defense against RNA viruses and they are effective against viruses that lack defense mechanisms such as the HIV-1-encoded *vif* (32).

To remove potentially inflammatory factors from the original HIV genome and improve overall vector safety, LVVs lack *vif*, rendering them defenseless to APOBEC3 family activity. It has been shown that, in *vif*-deficient HIV, APOBEC3G acted as a dominant inhibitor of viral replication causing either cDNA degradation or hypermutation, resulting in the inability to produce replication-competent virus (15). Recent reports examining APOBEC3 activity on LVVs show that the mutation rate can be reduced by 91.2% when viral vectors are packaged in an APOBEC3 knockout cell line (10). APOBEC3B has been shown to be carried over within LVV particles from incorporation during viral production to CAR T cells post-infection (33). Further, LVVs made in producer cells that overexpressed APOBEC3B resulted in lower overall CAR activity, whereas LVVs produced in APOBEC3B knockout producer cells was significantly associated with tumor cell death (e.g., robust CAR activity) post CAR T cell treatment (33). These data align with the observations presented here and call for the CAR T cell field to reassess current standards of LVV production practice. Specifically, both the third- and fourth-generation LVVs are produced in 293 or 293T fibroblast cell lines, which are known to actively express APOBEC3 genes. Together, the LVV SMRTcap data alongside those highlighted above support using APOBEC3 null cell lines for LVV production. If these alternatives are not scalable, the re-incorporation of HIV *vif* (or a variant thereof) into the LVV system could also be considered to block APOBEC3 hypermutation activity.

In conclusion, LVV SMRTcap provides a practical framework for higher-resolution QC of LVV-transduced cell products. Our results, together with other recent reports, highlight the impact of APOBEC3 activity on LVV mutagenesis and potential for reduced CAR T efficacy (10). Together these studies should motivate renewed attention to vector production conditions. Potential strategies to address this include broad and scalable utilization of producer cell lines engineered to reduce APOBEC3 activity, and vector design choices that reduce opportunities for RT- and recombination-driven deletions. More broadly, routine assessment of proviral integrity may help explain a portion of the product-to-product heterogeneity that contributes to variable clinical responses. LVV SMRTcap makes it feasible to measure this previously hidden layer of variation directly in research and clinical CAR T cell samples.

## Materials and methods

### Plasmid propagation and lentiviral production for research-derived CAR T cells

One Shot™ Stbl3™ chemically competent *E. coli* (Thermo Fisher Scientific, Irwindale, CA, USA, catalog no. C737303) plated on agar plates supplemented by ampicillin were obtained from Dr. William Tse and his team (University of Louisville, KY, USA). Colonies contained a CAR-carrying lentiviral plasmid derived from the FUGW vector (Addgene, Watertown, MA, USA, plasmid catalog no. 14883). *E. coli* carrying CAR-encoded LVV vector plasmids were expanded in LB broth, and plasmids were isolated using standard Maxi Prep procedures (Qiagen, Hilden, Germany, catalog no. 12162). Plasmids were confirmed to be the expected sequence by quality control sequencing using Oxford Nanopore (Plasmidsaurus, Louisville, KY, USA).

7mg of plasmid was diluted in 600uL of water for VSGV Lenti-X Packaging Single Shot (Takara Bio Inc., Shiga, Japan, catalog no. 631275) following the manufacturer’s instructions as written for transfection of Lenti-X HEK293T (Takara Bio Inc., catalog no. 632180) cultured in 85.5% DMEM (Millipore Sigma, Burlington, MA, USA, catalog no. D5796) supplemented with 10% heat inactivated tetracycline-free fetal bovine serum (Thermo Fisher Scientific, catalog no. A4736101), 1% Penicillin/Streptomycin (Thermo Fisher Scientific, catalog no. 15140-122), 1% Sodium Pyruvate Solution (Millipore Sigma, catalog no. S8636), 2.5% HEPES (Thermo Fisher Scientific, catalog no. 15630-080). Supernatant was collected at 48- and 72-hours post-transfection and collections were pooled into a single viral stock. Cell debris was separated from viral stock by centrifugation (500g for 10 minutes). Virus in clarified supernatant was then concentrated with the Lenti-X Concentrator (Takara Bio Inc., catalog no. 631231) and presence of viral particles was confirmed using Lenti-X GoStix Plus (Takara Bio Inc., catalog no. 631280). LVVs were stored at −80℃.

### Sequencing of viral stock RNA to investigate sources of internal deletions

Virus was purified by pelleting viral particles over a 20% sucrose gradient (Millipore Sigma, catalog no. S0389) by spinning at 21500 g for 2 hours at 4°C. Once pelleted, supernatant was removed. The viral pellet was resuspended in 120mL of phosphate-buffered saline (PBS) and allowed to rest on ice for one hour to loosen before resuspension. Viral mRNA was purified with the QIAamp Viral RNA Mini Kit (Qiagen, catalog no. 52904) according to manufacturer’s instructions. cDNA synthesis and amplification were performed using the NEBNext Single Cell/Low Input cDNA Synthesis & Amplification protocol (New England Biolabs, Inc., Ipswich, MA, USA, catalog no. E6421S). Amplified cDNA was prepped for sequencing using the SMRTbell prep kit 3.0 protocol REV06 (Pacific Biosciences, San Diego, CA, USA, catalog no. 102-141-700). SMRTbell library was annealed to sequencing primer and then bound to polymerase (SPRQ) prior to data collection on the Revio system using 30-hour movies. HiFi reads, defined as circular consensus sequences with empirical sequencing accuracy of Q30 (99.9%) or above were generated on system and used for all downstream analyses.

### LVV transduction of primary donor PBMC to generate “research” CAR T cells

Cryopreserved human healthy donor PBMCs were obtained from STEMCELL Technologies (Vancouver, Canada, catalog no. 70025.1). A 1-2×10^6^ cells/mL cell density was maintained in 85.5% Roswell Park Memorial Institute (RPMI) media, 10% Heat Inactivated fetal bovine serum (FBS), 1% PenStrep, 1% mL GlutaMax (Thermo Fisher Scientific, catalog no. 35050-061), 2.5% N-2-hydroxyethylpiperazine-N-2-ethane sulfonic acid (HEPES). Cytokines IL-21 (Thermo Fisher Scientific, catalog no. 200-21-10UG), IL-4 (Thermo Fisher Scientific, catalog no. 200-04-20UG), and IL-7 (Thermo Fisher Scientific, catalog no. 200-07-10UG) each were added into the media at a concentration of at 20 ng/mL upon plating. PBMCs were activated with Dynabeads Human T-Activator CD3/CD28 (Thermo Fisher Scientific, catalog no. 11132D) in a 1:1 ratio cells-to-beads for 72hr and removed. PBMCs were incubated with LVV stock (1×10^6^ cells in 300mL concentrated viral stock) and incubated for 4 hours at 37°C+5% CO_2_. Media was added to dilute the infection and cells (at 1mL media for every million cells) continue to incubate with the virus in 37°C+5% CO_2_ for 48h. Virus was removed from PBMCs by pelleting cells via centrifugation at 500g for 10 minutes. PBMCs were washed 3 times using PBS with 10% FBS and recovered overnight in media. gDNA and RNA from LVV-transduced CAR T cells were co-extracted using the All Prep kit (Qiagen, catalog no. 80204) for downstream LVV SMRTcap data analysis.

### Generation and collection of clinically derived CAR T cells

Clinical PBMC samples were obtained longitudinally from a participant with multiple myeloma enrolled on the ongoing clinical trial NCT03549442. Samples analyzed in this study were collected on Day +7, +14, +21, and +28 following CAR T cell infusion. Patient CAR T cell product generation, collection, and clinical administration were performed as previously described by Cohen et al (34). The study was conducted under a University of Pennsylvania IRB-approved clinical protocol (IRB# 822756), and samples were obtained after written informed consent in accordance with the Declaration of Helsinki.

### LM-PCR Preparation

Genomic DNA was fragmented to ∼500bp by sonication (three rounds: 4x 5 sec with 10 sec rest at 20 amplitude). gDNA was end-repaired using the Anza DNA Blunt End Kit (Thermo Fisher Scientific, catalog no. IVGN2404), 1 µl of DNA Blunting Enzyme Mix, 2 µl of 10X Blunting Buffer, and 1 µg of fragmented DNA were mixed and brought to a final volume of 20 µl with nuclease-free water. This reaction was incubated at 20°C for 15 minutes and purified using AMPure PB beads (Pacific Biosciences, catalog no. 100-265-900) at a vol:vol ratio of 1.2X beads-to-reaction. Following purification, the fragmented DNA was then A-tailed using 5µL of 10X NEBuffer 2, 0.5 µL of 10 mM dATP, 3 µL of Klenow Fragment (New England Biolabs, Inc., catalog no. M0212S), and purified DNA, with nuclease-free water added to obtain a final volume of 50 µl. The mixture was incubated at 37°C for 30 minutes and purified using AMPure PB beads (Pacific Biosciences, catalog no. 100-265-900) at a vol:vol ratio of 1.2X beads-to-reaction.

Linkers were prepared by annealing 3.5 µL each of the short and long oligonucleotide strands (100 µM in 10 mM Tris-HCl, pH 8.0, 0.1 mM EDTA) with 28 µl of 10 mM Tris-HCl, pH 8.0, 0.1 mM EDTA and heated to 90°C and cooled to 20°C at a rate of 1°C per minute. Annealed linkers were ligated to sheared, A-tailed and purified gDNA using 7.5 µL of annealed linker, 1 µg DNA, 2 µL of T4 DNA Ligase (800 units) (New England Biolabs, Inc., catalog no. M0202S), and 2 µL of T4 DNA Ligase Reaction Buffer, with nuclease-free water added to achieve a final volume of 50 µL, followed by incubation overnight at 12°C and then purified using AMPure PB beads (Pacific Biosciences, catalog no. 100-265-900) at a vol:vol ratio of 1.2X beads-to-reaction. Linker-ligated gDNA was digested with 1 µL of BglII (100 units) (New England Biolabs, Inc., catalog no. R0144S), 1 µg of DNA, and 5 µL of 10X NEBuffer, with nuclease-free water added to achieve a final volume of 50 µl. The mixture was incubated at 37°C for 2 hours and purified then purified using AMPure PB beads (Pacific Biosciences, catalog no. 100-265-900) at a vol:vol ratio of 1.2X beads-to-reaction.

The first-round PCR was performed in 4 parallel reactions each including: 2.5 µL of 15 µM LTR primer, 0.5 µL of 15 µM linker-specific primer, 2.5 µL of 10X PCR buffer, 0.5 µL of 2.5 mM dNTPs, 0.5 µL of DNA Advantage 2 Polymerase (Takara Bio Inc., catalog no. 639201), and 100 ng of digested linker-ligated DNA, with nuclease-free water added to obtain a final volume of 25 µL. First round thermocycling conditions were: initial denaturation at 94°C for 2 minutes; 30 cycles of denaturation at 94°C for 15 seconds, annealing at 55°C for 30 seconds, and extension at 68°C for 45 seconds; followed by a final extension at 68°C for 10 minutes. PCR products were purified and prepared for a second-round of PCR. PCR 2 was performed identical to the first PCR, however the first-round LTR primer was replaced with the second-round LTR primer that contains a barcode sequence used for pooling samples and specified by Serrao et al. (27). Final PCR products were pooled, and sequenced as a 2×250 on an Illumina MiSeq V3 flow cell.

### LM-PCR analysis

Only R1 LM-PCR reads were used and mapped to the hg38 genome reference since most of the sequencing fragments were smaller than 250 bp. Sequences were mapped using blat v 37×1 (35) with simple repeat masking turned off. Reads mapping to multiple locations were filtered out using blat’s pslReps functionality. Locations were filtered for those that had at least one hit, at least 10 hits, at least 100 hits, and at least 1000 hits. We focused on those integration locations with at least 10 hits to reduce potential false positive locations (Figure 1C). Given that no unique molecular identifiers (UMIs) were used in the sequencing and LM-PCR amplifies based on a common set of primers, we were unable to distinguish between PCR replicates and clonal expansion events.

### LVV SMRTcap

The LVV-SMRTcap protocol used here closely follows that previously shown (26). Briefly, gDNA was fragmented to ∼10kb by g-TUBE (Covaris, Woburn, MA, USA, catalog no. 520079) centrifuging at 4100 rpm using an Eppendorf 5242R microcentrifuge. Sheared gDNA was purified and concentrated with AMPure PB beads (Pacific Biosciences, catalog no. 100-265-900) using a 0.5X beads-to-sample vol:vol ratio. The Long Read Library Preparation and Standard Hyb v2 Enrichment protocol (Twist Bioscience, San Francisco, CA, USA) was followed according to the manufacturer’s guidelines with some modifications. Sheared gDNA was end repaired and A-tailed. Barcoded adapter oligo pairs (36) were ligated to the ends of DNA fragments, and purified using AMPure PB beads (Pacific Biosciences, catalog no. 100-265-900) using a 0.5X beads-to-sample vol:vol ratio. Samples were mass amplified using PrimeSTAR GXL Polymerase (Takara Bio Inc., catalog no. R050B), and primer binding regions in barcoded adapters. Fragment size and mass was measured using TapeStation 4150 (Agilent, Santa Clara, CA, USA) and Qubit fluorometer 4 (Thermo Fisher Scientific). Fragments >8kb were isolated using the BluePippin system (Sage Science, Beverly, MA, USA), and fragment size and mass was measured again.

Barcoded libraries were equimolar pooled and enriched by desiccation in a vacuum concentrator and reconstituted in a small volume of Blocker Solution (Twist Biosciences, catalog no. 100774) and mixed with Hybridization Mix (Twist Biosciences, catalog no. 100528) in combination with a custom probe pool designed for LVV capture. Final mix was covered with the Hybridization Enhancer (Twist Biosciences, catalog no. 100986) and hybridized at 70°C for 16–18 hours. Fragments were isolated by incubating the pool with M-270 Dynabeads (Invitrogen, Carlsbad, CA, USA, catalog no. 65305) and mixing gently end-over-end for 30 minutes at room temperature. Multiple wash steps with Wash Buffer I (Twist Biosciences, catalog no. 100589) and II and (Twist Biosciences, catalog no. 100590) removed fragments not captured, including low-affinity, off-target fragments. Fragments were eluted from beads using a mixture of water, 0.2N NaOH, and 200 mM Tris-HCl. Enriched material was amplified using the KOD Xtreme Hot Start Polymerase (Millipore Sigma, catalog no. 71975-M) in replicates of two, as recommended by the manufacturer. Fragment size and mass were measured. Samples were prepped for sequencing using the SMRTbell prep kit 3.0 protocol REV06 (Pacific Biosciences, catalog no. 102-141-700). SMRTbell library was annealed to the Revio sequencing primer, followed by polymerase (SPRQ) binding. Data was collected on the Revio system using 30hr movies. Following data collection, circular consensus sequences with empirical sequencing accuracy of Q30 (99.9%) or above were generated on Revio system, output as “HiFi reads,” and used for downstream custom analyses described below.

### LVV SMRTcap analyses

All data were analyzed by a three-step pipeline to: (i) identify the sequencing reads that contained LVV content; (ii) define integration sites and enumerate expanded clones; and (iii) characterize proviral integrity per molecule. Details for each step are provided below.

### Identifying LVV integrations and host flanking sequences

LVV SMRTcap HiFi reads were mapped to the LVV plasmid sequence using minimap2 v2.24-r1122 with the parameters -t 16 -m 0 -Y -ax map-pb to identify reads with LVV integrations (37). Reads were then filtered to exclude secondary alignments and unmapped reads using samtools view v1.16.1 with the parameters -h - S -F 260 (38). Reads were further filtered to exclude secondary alignments (-F 256), unmapped reads (-F 4) and supplementary alignments (-F 2048) using the samtools view command with the -F 2308 parameter. Additional parameters -h and -S were included to retain the header and specify the input format as SAM. The filtered reads were then hard masked for the regions containing hits to the LVV reference and iteratively searched again using minimap2 with the parameters -t 16 -Y -p 0 -N 10000 -ax map-pb until no additional LVV hits were found which allows for the detection of duplicates and rearrangements. At the end of this iteration, all regions of the reads containing an LVV hit were hard masked. The hard masked reads were then searched against the primary GRCh38 human genome assembly to identify insertion site flanks using minimap2 and the parameters -t 16 -Y -p 0 -N 10000 -ax map-pb which were then filtered to remove supplementary alignments, secondary alignments, and unmapped reads with samtools view and the parameters -h -S -F 2308. The resulting reads were then processed using a custom python script that parses the sam alignment files to produce a tab-delimited file containing information about the LVV sequence hit, and the left and right human flanking sequences.

### Mapping human flanking sequences and assigning clonal expansion events

The LVV-mapped summary.csv file was imported into the R (version 4.3.1) pipeline via openxlsx (version 4.2.6.1). Data was cleaned through use of tidyr (version 1.3.1), dplyr (version 1.1.4), purrr (version 1.02), stringr (version 1.5.1), and stringi (version 1.8.4) (39–43). Quality control of reads was performed through examination of human genome flanking coordinates to identify mis-mapped reads or paired integration sites existing at disparate genomic loci (e.g., 5’ end on chr2 with 3’ end on chr17), which were interpreted as indicative of PCR artifact. Reads that did not pass these quality parameters were removed from further analyses. Clonal expansion events were defined as those molecules that contained identical integration sites, but unique shearing end coordinates (schematized in Figure 4A). Integration sites were considered to have high confidence mapping if they were within a 10Dbp window of each other (± 5Dbp up- or downstream) to account for polymerase slippage during amplification. 10Dbp was chosen as the maximal spread with which the 5’ and 3’ integration sites could differ because of the well-described polymerase slippage of up to 5Dbp and existing technical difficulty in many sequencing platforms with homopolymeric sequence identity (44, 45). Similarly, integration coordinates between different reads that overlapped (e.g., chr1:300,600,428–300,600,432 and chr1:300,600,430–300,600,433) were considered clones of the same initial LVV integration event with discrepancies assumed to be from the sequencing process. In parallel, PCR duplications were identified as molecules that shared both integration sites and shearing coordinates. The coordinate, clonal expansion, and PCR data were mapped to genes (including identification of sub-genomic features), intergenic space, repeat elements, and ENCODE features (e.g., CpG Islands, promoters, enhancers) as pulled from the UCSC Genome Browser via BiocManager (version 1.30.24), GenomicRanges (version 1.52.1) and plyranges (1.20.0). All associated R software package dependencies were used with default settings.

### Gene intactness

A custom perl script findCARTGeneRegions was used to determine the intactness of the overall integration. This script measured the intactness of the UbC promoter in Figure 2D. This was done by comparing the LVV integration sequence against the respective annotated region from the closest specified LVV reference genome identified during the mapping step using NCBI blastn v2.10.0+ (46) with the parameter - word_size 16 to remove spurious hits. If no hits were found for a particular region, that region was assigned a value of 0. If the length and identity of the match was between 0 and 50% of the reference region, then that gene was assigned a value of 1. Identity between 50-95% was valued at 2. Length and sequence identity matches greater than 95% were assigned a value of 3 (identity of 95% and above qualified as an “intact” call). Post-processing of the integration strings was performed to change 0 values at the end to “-“ if host flanking sequences were not available, indicating a loss of content to mechanical shearing.

### Proviral integrity visualization

Fastq files were aligned to reference fasta using minimap2 for the downstream creation of BAM files needed for data visualization. Proviral visualization was done using Integrative Genomics Viewer (IGV) version 2.17.2 (47). Visualization was filtered to show insertions >2bp. Read histograms indicate SNV positions if a SNV is in at least 10% of reads.

## Data Availability Statement

All sequencing data described here, including total HiFi reads per sample and the post-filtered total LVV-containing reads per sample, have been deposited into the Sequence Read Archive and can be retrieved from BioProject (ONGOING). Custom computational tools are available through GitHub at URL: (ONGOING).

## Acknowledgments

Support provided by NIH grants F31 AI186545 (Kaiser) and P20GM103436 (Rouchka). SMRT sequencing instrumentation support by NIH grant S10OD034432-01 (Smith). Contents of this work are the sole responsibility of the authors and do not reflect official positions of the National Institutes of Health. The authors want to extend deep gratitude to the University of Louisville Sequencing Technology Center staff for close partnership that ensured robust data production for this new, off-specification long-read protocol. The authors would like to thank Dr. William Tse for providing materials and subject matter expertise. The authors would like to thank Dr. Kamille Rasche for editorial and figure feedback. “SMRT” is a registered trademark of Pacific Biosciences, Inc. The authors would like to thank the patients who contributed the samples studied here.

## Author Contributions

**Catherine W. Kaiser**

**Roles** Conceptualization, Data curation, Formal analysis, Investigation, Methodology, Project administration, Resources, Software, Supervision, Validation, Visualization, Writing – original draft, Writing – review & editing

**Ghazal S. Mehs**

**Roles** Data curation, Methodology, Review & editing

**Erin M. Elliott**

**Roles** Data curation, Methodology, Review & editing

**Joanna E. Mroczkowska**

**Roles** Methodology, Resources, Review & editing

**Michael Ferguson**

**Roles** Methodology, Resources, Review & editing

**Alfred L. Garfall**

**Roles** Resources

**Frederic D. Bushman**

**Roles** Conceptualization, Writing – review & editing

**Joseph A. Frietta**

**Roles** Resources, Review & editing

**Eric C. Rouchka**

**Roles** Conceptualization, Data curation, Formal analysis, Funding acquisition, Software, Supervision, Validation, Writing – original draft, Writing – review & editing

**Melissa L. Smith**

**Roles** Conceptualization, Data curation, Formal analysis, Funding acquisition, Project administration, Supervision, Writing – original draft, Writing – review & editing

