## Supplemental Materials for "LVV SMRTcap reveals extensive proviral variation in lentiviral vector-transduced CAR T cells"

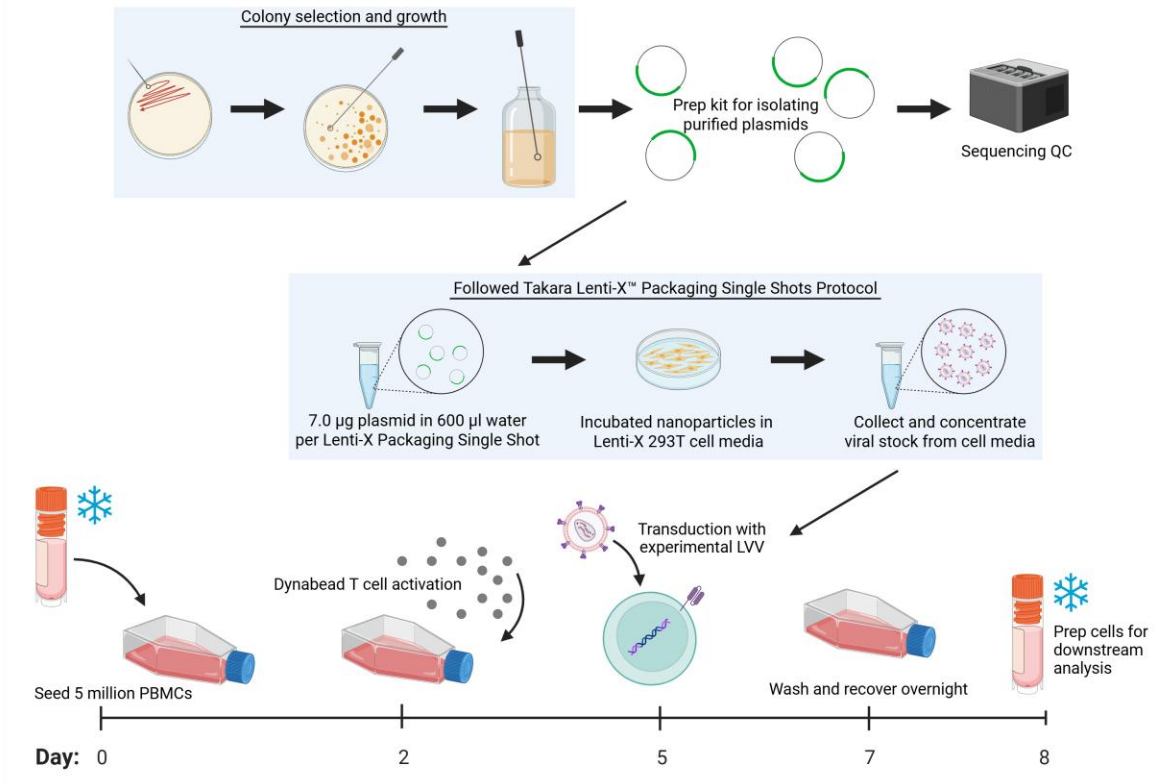


**Figure S1: Experimental layout of research-derived CAR T cell preparation.** *E.coli* was plated on agar that was used for isolating viral plasmid. Plasmid underwent QC sequencing, and 7ug was used for viral production with the Takara Lenti-X Packaging Single Shot System (Takara Bio Inc.). Five million PBMCs were plated and activated prior to transduction. PBMCs were incubated with virus for 48 hours at which point the virus was washed off and cells were allowed to recover overnight and then were stored in liquid nitrogen. Created in BioRender.com.

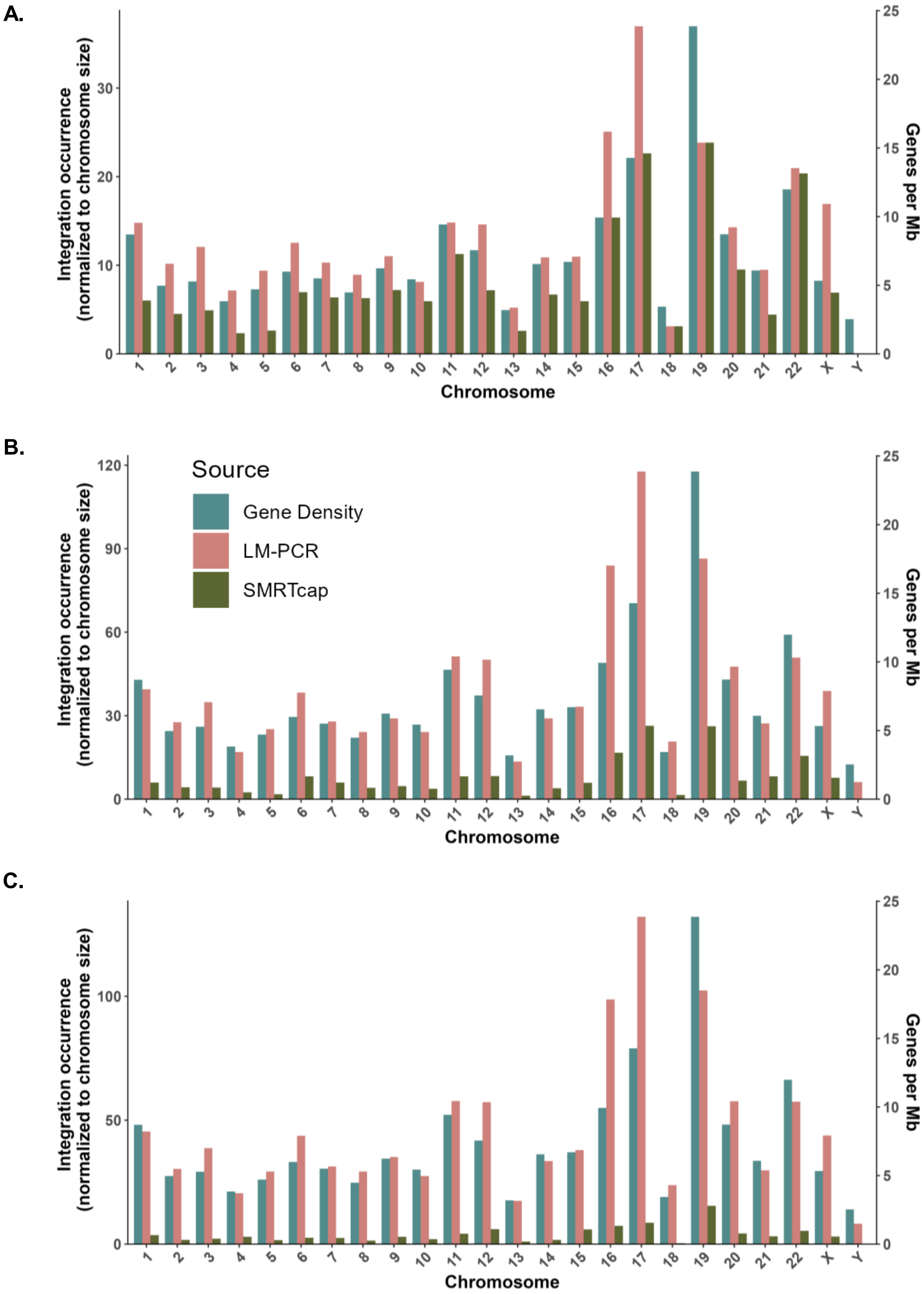


**Figure S2: LVV SMRTcap and LM-PCR viral integration coverage per chromosome from donors A) 1001, B) 5002, C) 4006.** Number of integrations are normalized to chromosome size. Gene density is presented using Cold Spring Harbor Laboratory Press's Guide to the Human Genome, genes per sequenced Mb is recorded for every chromosome (47).

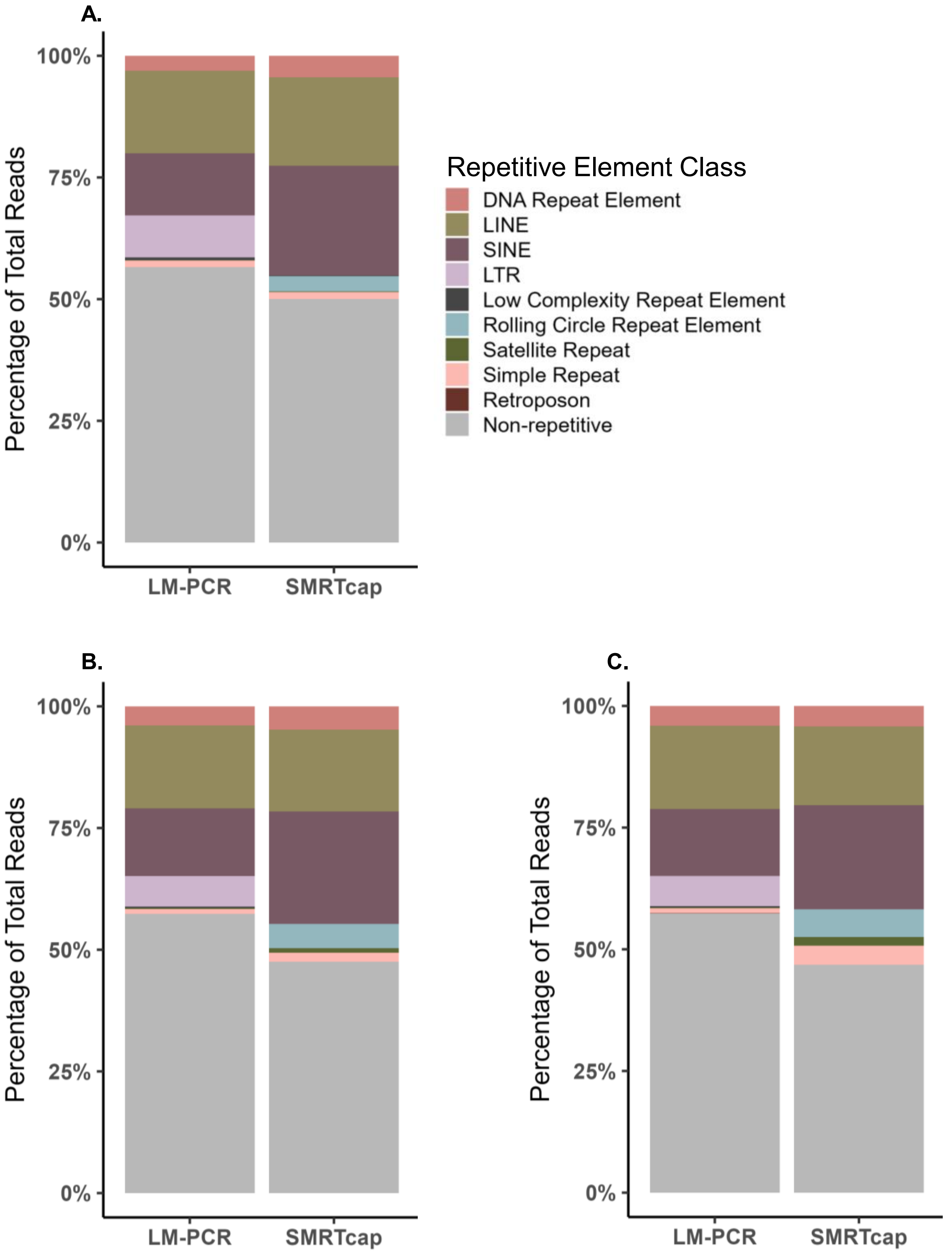


**Figure S3: LVV SMRTcap and LM-PCR integration site composition into unique and repetitive elements from donors A) 1001, B) 5002, C) 4006.**

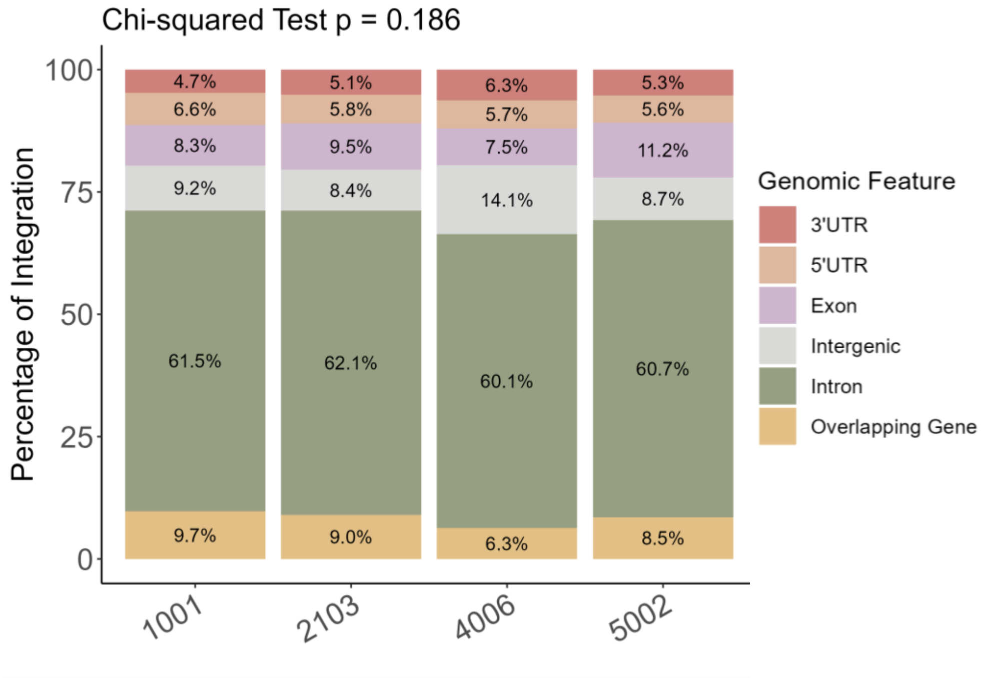


**Figure S4: Genomic features of integration in LVV SMRTcap reads for research-grade CAR T cells.** Chi-squared test compares the distribution of genomic integration features across CAR T cell samples.

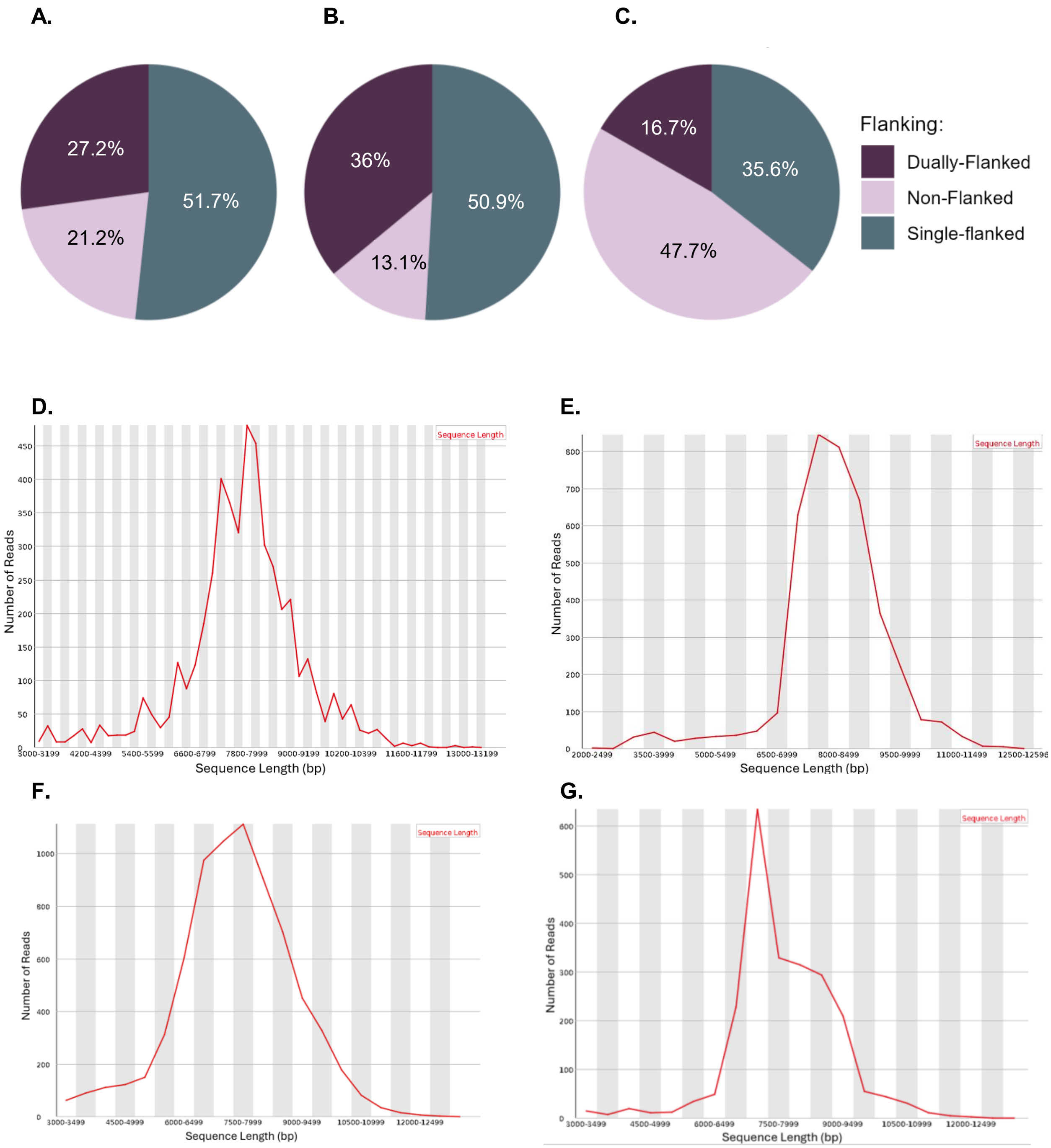


**Figure S5: LVV SMRTcap shearing profiles for research-grade CAR T cells.** Viral shearing read length profiles for donors **A)** 1001, **B)** 5002, **C)** 4006. SMRTcap sequence read length profiles for donors **D)** 1001, **E)** 5002, **F)** 2103, and **G)** 4006.

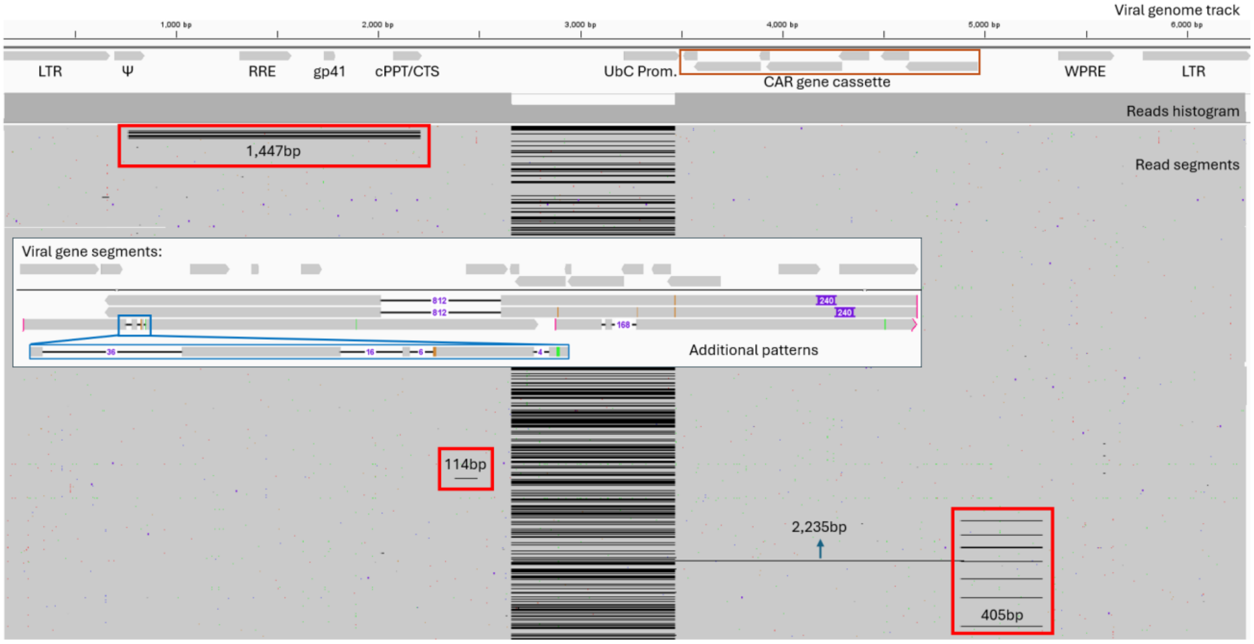


**Figure S6: Characterized indels in research CAR T cells for donor 1001 using LVV SMRTcap.** Dually flanked proviruses mapped back to the viral genome reference and visualized using IGV. Viral genes are annotated beneath the viral genome track. Read segments include deletions, annotated as black lines. An expanded view of read segments shows additional deletion patterns found in single-flanked reads.

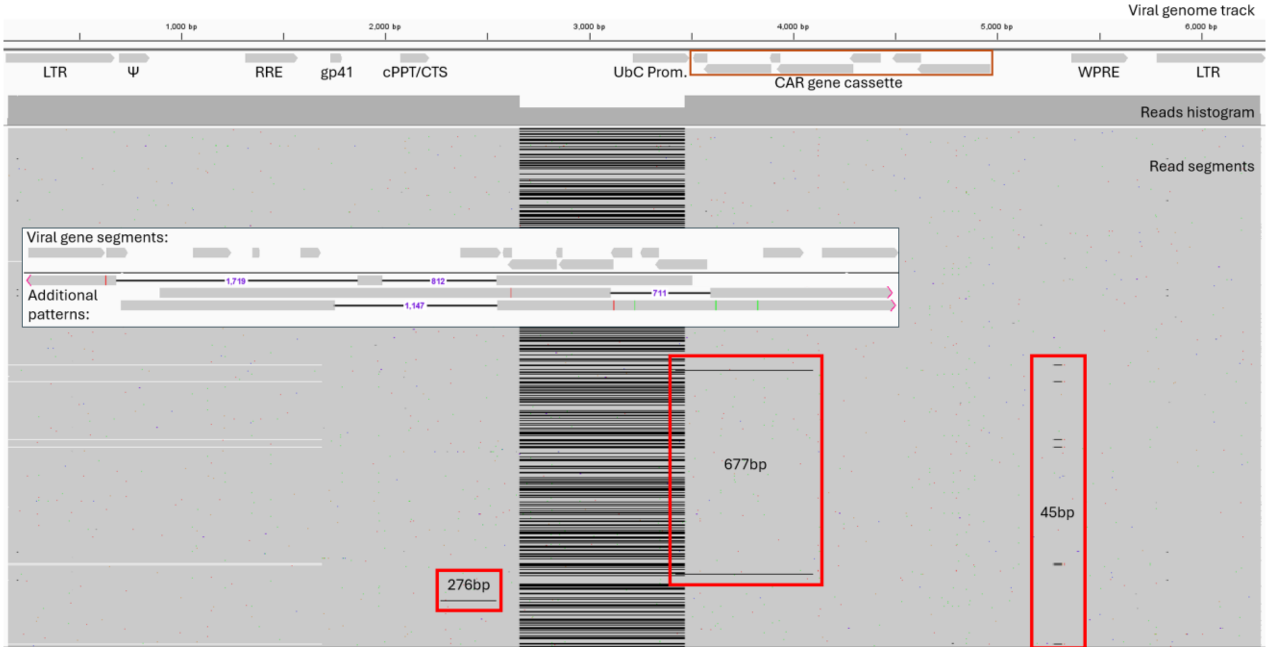


**Figure S7: Characterized deletions in research CAR T cells for donor 5002 using LVV SMRTcap.** Dually flanked proviruses mapped back to the viral genome reference and visualized using IGV. Viral genes are annotated beneath the viral genome track. Read segments include deletions, annotated as black lines. An expanded view of read segments shows additional deletion patterns found in single-flanked reads.

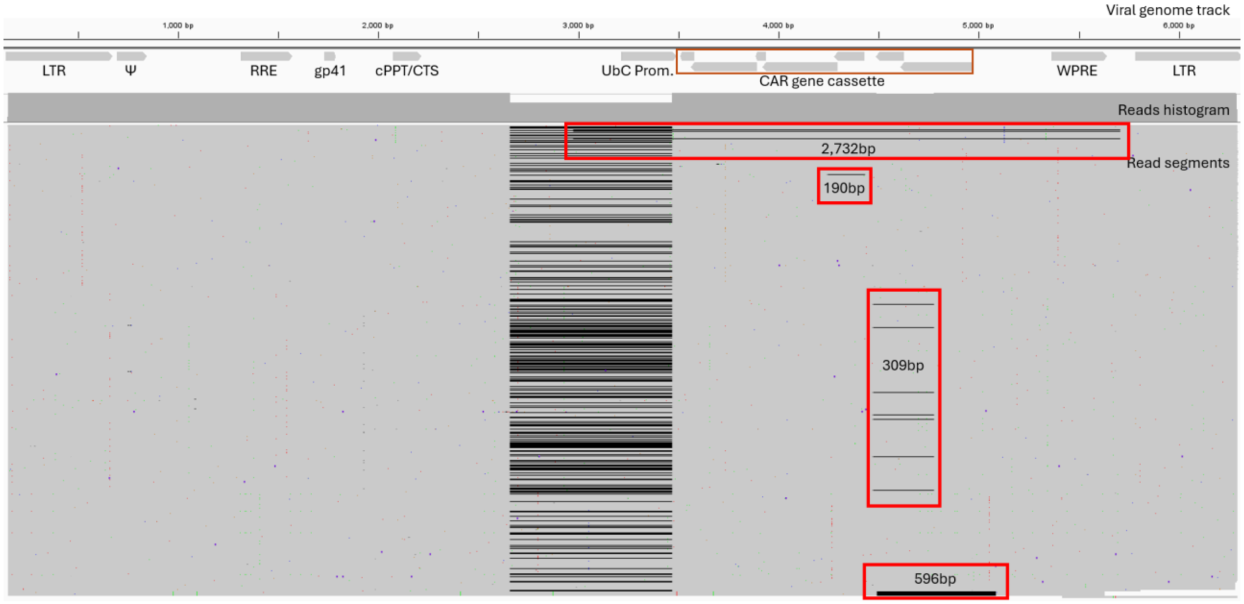


**Figure S8: Characterized deletions in research CAR T cells for donor 4006 using LVV SMRTcap.** Dually flanked proviruses mapped back to the viral genome reference and visualized using IGV. Viral genes are annotated beneath the viral genome track. Read segments include deletions, annotated as black lines.

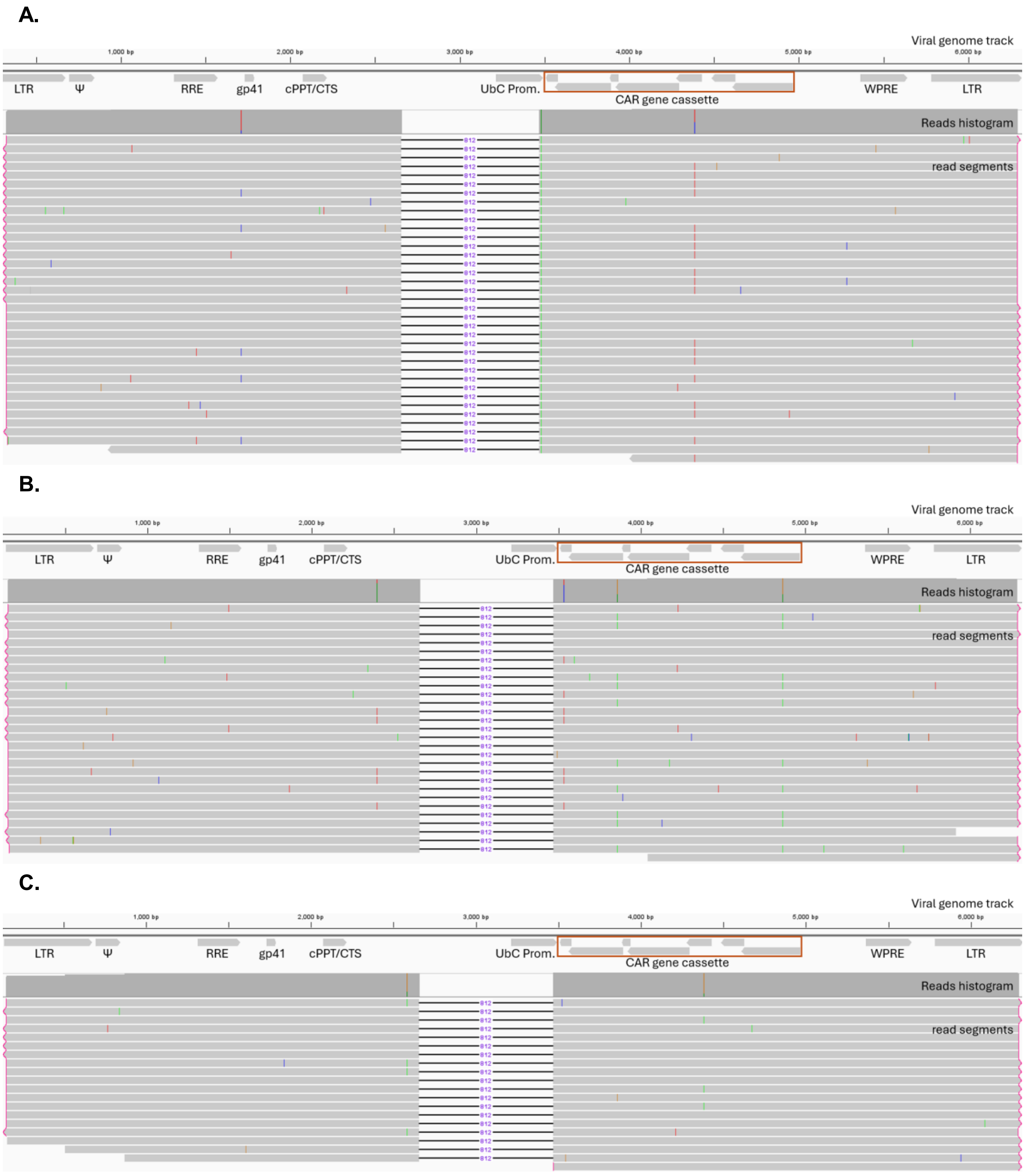


**Figure S9: Clonally expanded LVV integration for research-grade CAR T cells for donors A) 1001 integrated into *TBCD*, B) 5002 integrated into *ENSG00000261499*, and C) 4006 integrated into *OTUD4*.** Varying colors represent unique DNA bases, and highlight SNPs: A = green, G = orange, T = red, and C= blue. The read histogram shows SNPs occurring in >10% of reads.

**A.**

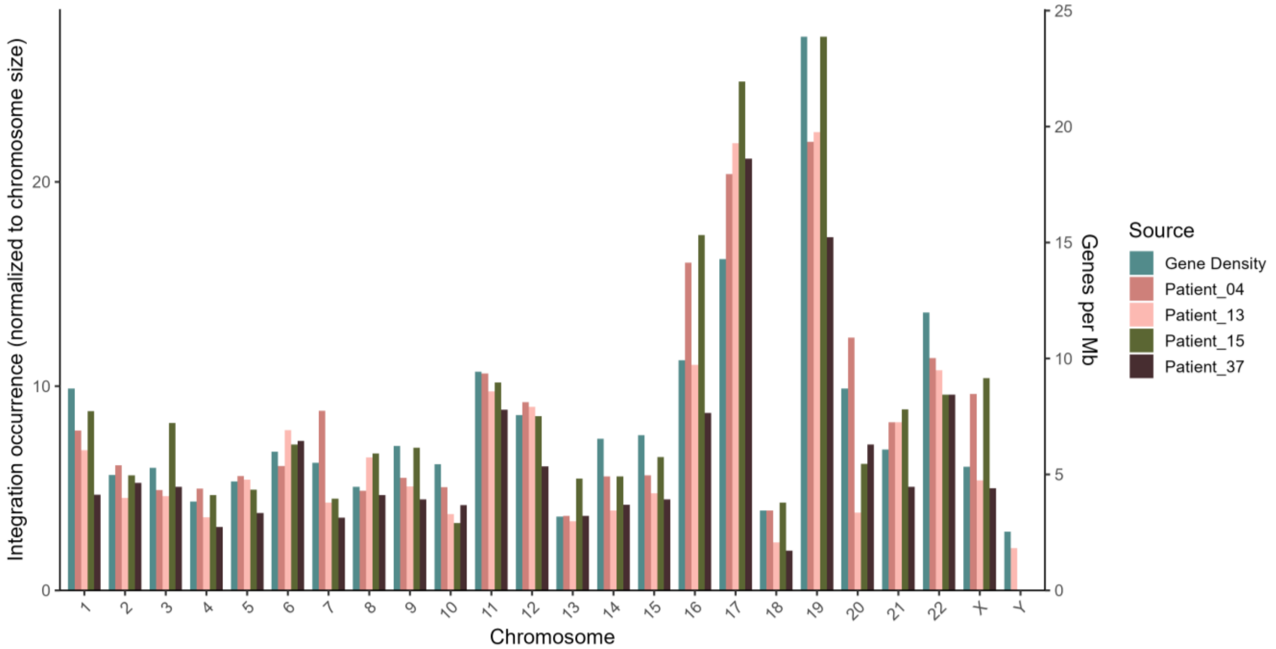


**B.**

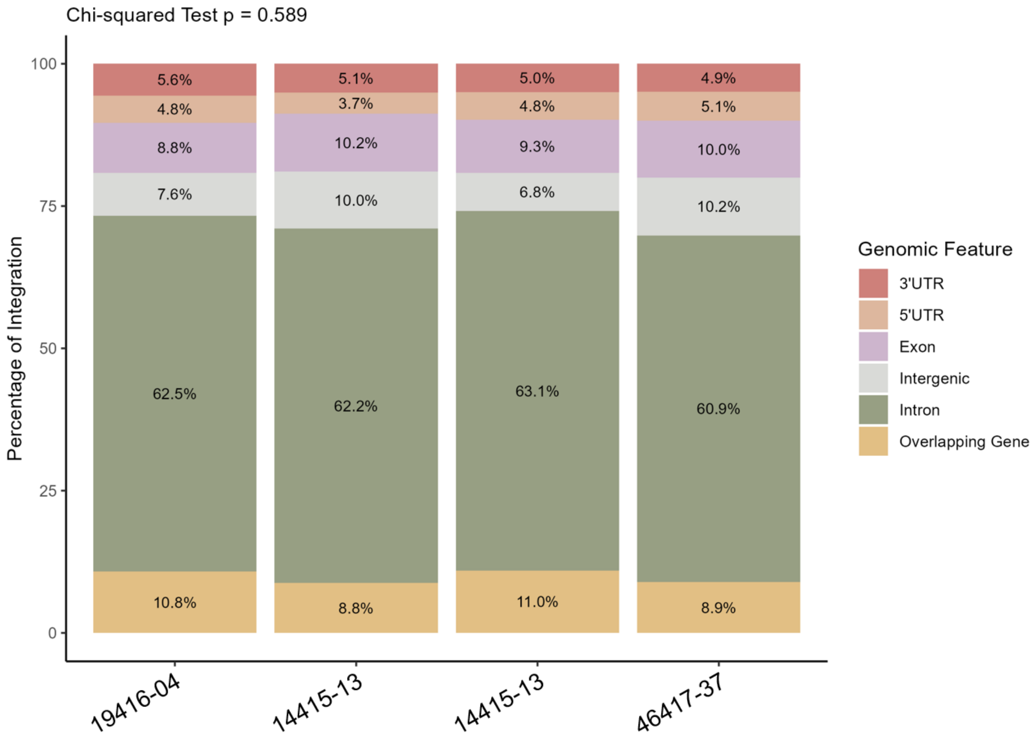


**Figure S10: LVV integration profiles for clinically derived CAR T cell products. A)** LVV SMRTcap and LM-PCR viral integration coverage per chromosome. Number of integrations are normalized to chromosome size. Gene density is presented using Cold Spring Harbor Laboratory Press's Guide to the Human Genome, genes per sequenced Mb is recorded for every chromosome (47). **B)** Genomic features of integration in SMRTcap reads. Chi-squared test compares the distribution of genomic integration features across CAR T cell samples.

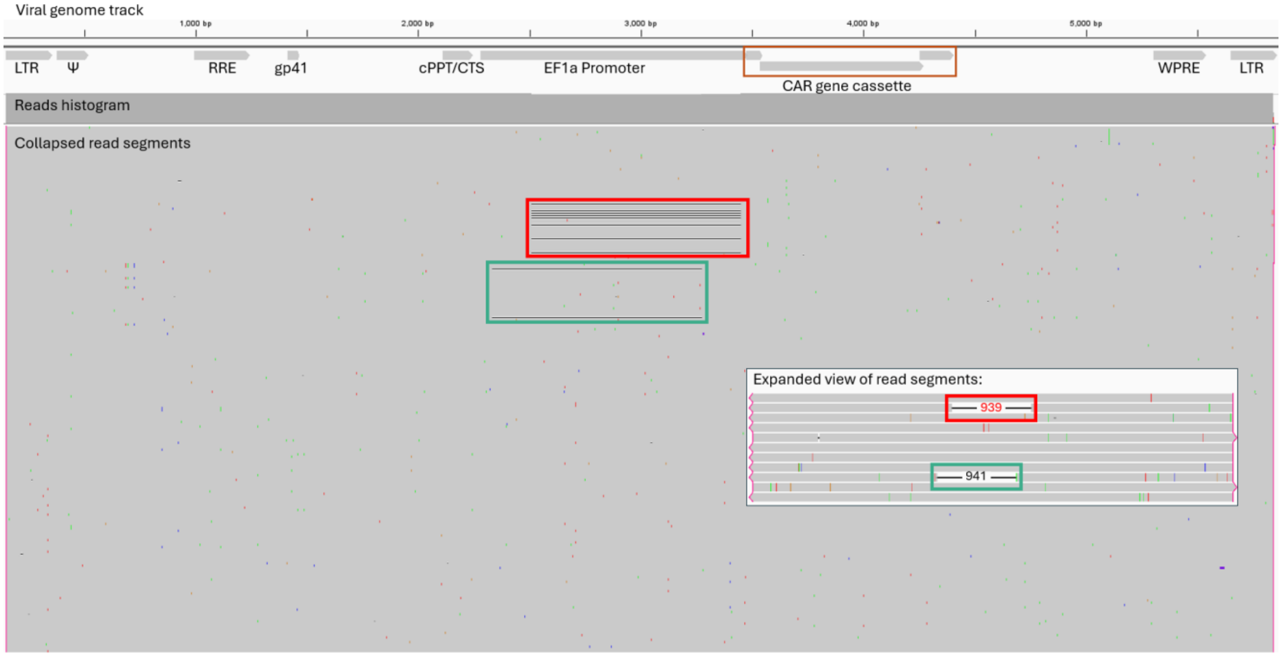


**Figure S11: Characterized deletions in clinically derived CAR T cells for patient 46417-37 using LVV SMRTcap.** Dually flanked proviruses mapped back to the viral genome reference and visualized using IGV. Viral genes are annotated beneath the viral genome track. Read segments include deletions, annotated as black lines. An expanded view of read segments shows recurring deletions with matching outlines to show their position in the unexpanded view.

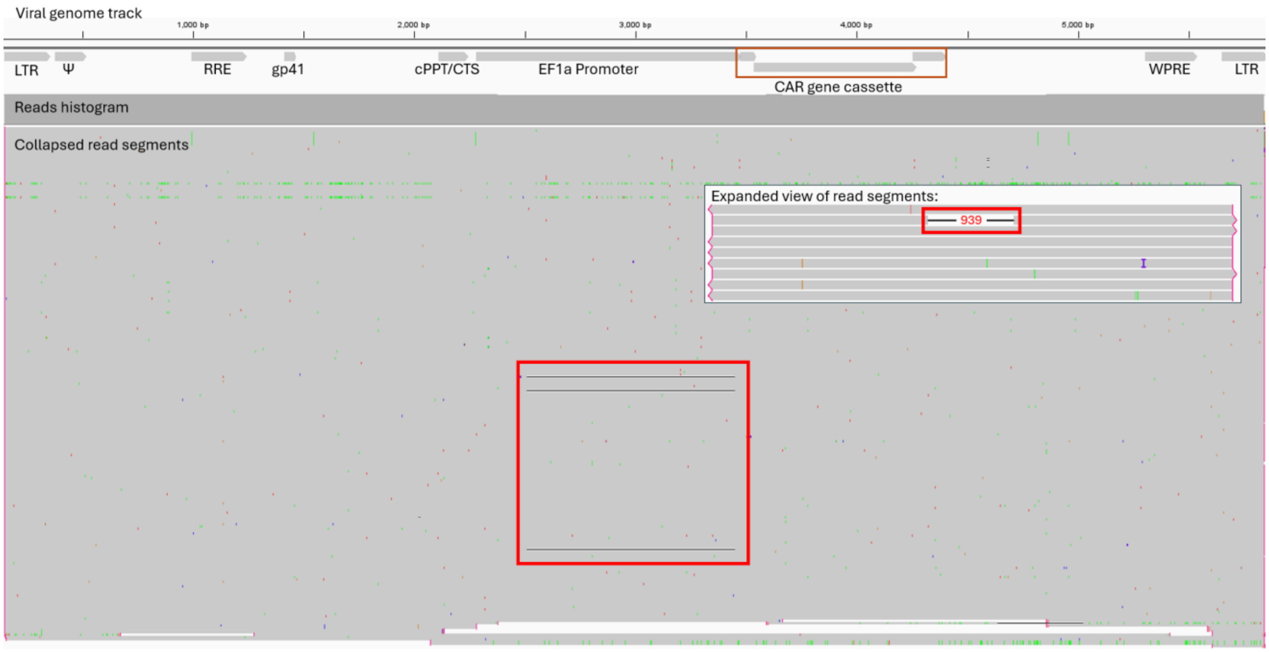


**Figure S12: Characterized deletions in clinically derived CAR T cells for patient 14415-13 using LVV SMRTcap.** Dually flanked proviruses mapped back to the viral genome reference and visualized using IGV. Viral genes are annotated beneath the viral genome track. Read segments include deletions, annotated as black lines. An expanded view of read segments shows recurring deletions with matching outlines to show their position in the unexpanded view.

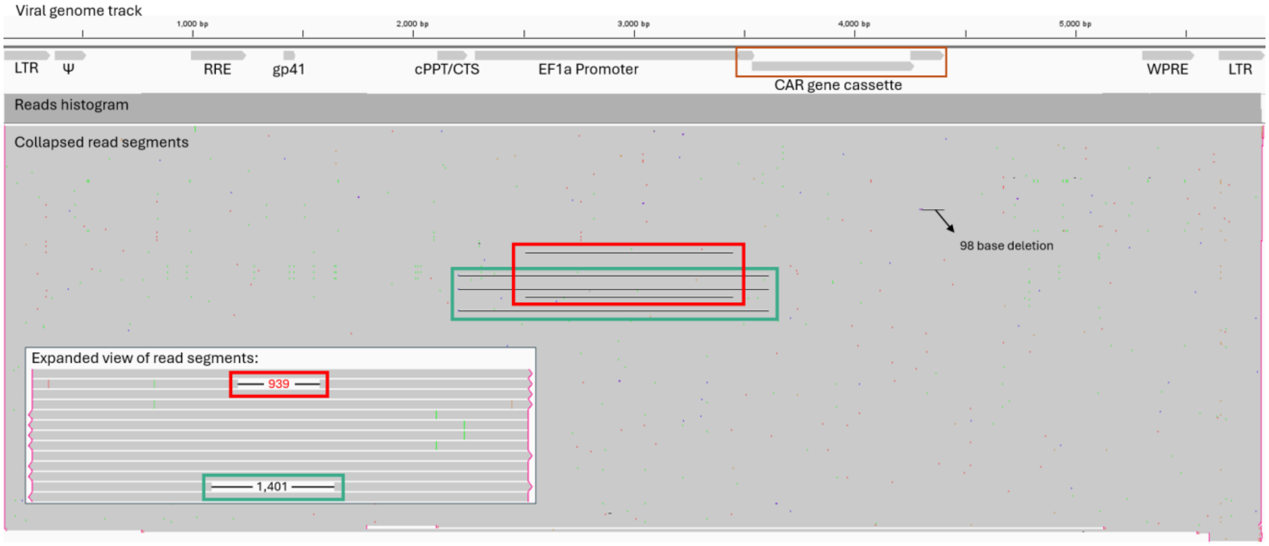


**Figure S13: Characterized deletions in clinically derived CAR T cells for patient 14415-15 using LVV SMRTcap.** Dually flanked proviruses mapped back to the viral genome reference and visualized using IGV. Viral genes are annotated beneath the viral genome track. Read segments include deletions, annotated as black lines. An expanded view of read segments shows recurring deletions with matching outlines to show their position in the unexpanded view.

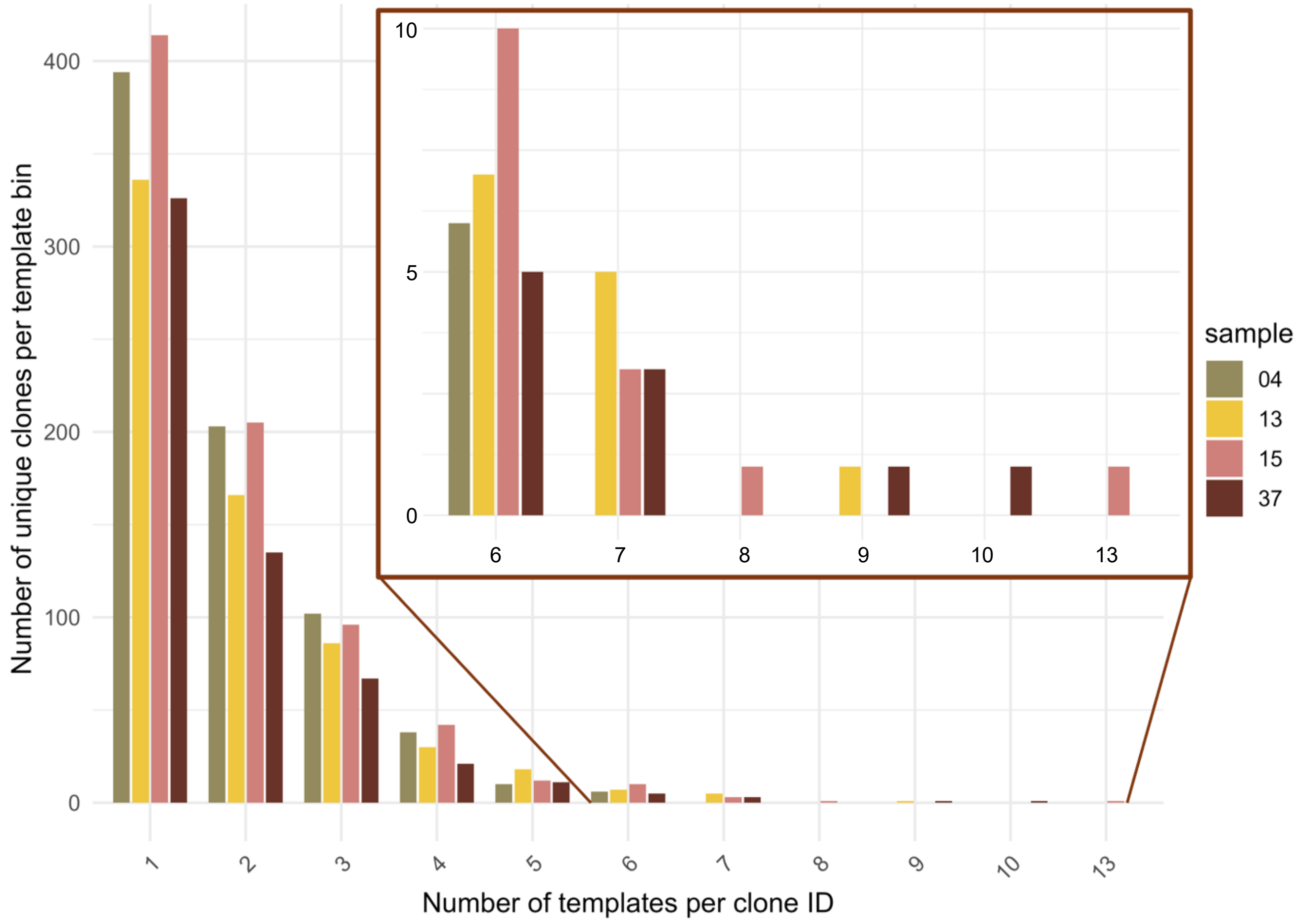


**Figure S14: Histogram of clonal expansion across clinical CAR T cell product samples.** The x-axis represents the number of templates for an individual clonal group, while the y-axis is the number of templates per clonal group; for example, x = 1 is a unique clone, whereas x = 3 is a clonal group containing 3 unique templates.
